# Autism and Intellectual Disability-Associated *MYT1L* Mutation Alters Human Cortical Interneuron Differentiation, Maturation, and Physiology

**DOI:** 10.1101/2024.09.11.612541

**Authors:** Ramachandran Prakasam, Julianna Determan, Mishka Narasimhan, Renata Shen, Maamoon Saleh, Gareth Chapman, Komal Kaushik, Paul Gontarz, Kesavan Meganathan, Bilal Hakim, Bo Zhang, James E. Huettner, Kristen L. Kroll

## Abstract

MYT1L is a neuronal transcription factor highly expressed in the developing and adult brain. While pathogenic *MYT1L* mutation causes neurodevelopmental disorders, these have not been characterized in human models of neurodevelopment. Here, we defined the consequences of pathogenic *MYT1L* mutation in human pluripotent stem cell-derived cortical interneurons. During differentiation, mutation reduced MYT1L expression and increased progenitor cell cycle exit and neuronal differentiation and synapse-related gene expression, morphological complexity, and synaptic puncta formation. Conversely, interneuron maturation was compromised, while variant neurons exhibited altered sodium and potassium channel activity and reduced function in electrophysiological analyses. CRISPRi-based knockdown similarly impaired interneuron differentiation and maturation, supporting loss of function-based effects. We further defined MYT1L genome-wide occupancy in interneurons and related this to the transcriptomic dysregulation resulting from *MYT1L* mutation, to identify direct targets that could mediate these phenotypic consequences. Together, this work delineates contributors to the etiology of neurodevelopmental disorders resulting from *MYT1L* mutation.

## INTRODUCTION

Intellectual and developmental disorders (IDDs), including autism spectrum disorder (ASD), intellectual disability (ID), and attention deficit hyperactivity disorder (ADHD) frequently co-occur in human subjects, with clinical manifestations including impaired social interaction, communication, and repetitive behaviors. In recent years, genetic liabilities have been defined, including *de novo* and inherited variants in several hundred genes (Gaugler et al. (2014); Havdahl et al. (2021); Zhou et al. (2022)). A fraction have been studied in human stem cell and/or animal models and often involve dysregulation of the balance between neuronal inhibition and excitation in the cerebral cortex, due to altered development of cortical excitatory neurons, inhibitory interneurons, or both cell types (Lewis et al. (2019); Meganathan et al. (2021); Kathuria et al. (2018)). Among these, pathogenic variants in the gene encoding the proneuronal transcription factor Myelin transcription factor 1 like (MYT1L) were recently defined as a new contributor to IDDs (De Rocker et al. (2015); Blanchet et al. (2017); Coursimault et al. (2022); Chen et al. (2021)).

MYT1L is a transcription factor in the neural-specific myelin transcription factor 1 family, zinc finger transcription factors that recognize a consensus DNA binding motif (Manukyan et al. (2018)). Initial work suggested a role in promoting neural development, as overexpression (OE) with two other neurogenic transcription factors converted mouse embryonic fibroblasts into functional neurons (Vierbuchen et al. (2010)), while MYT1L OE in P19 embryonic carcinoma cells increased neuronal differentiation (Mall et al. (2017); Romm et al. (2005)). Conversely, shRNA knockdown converted human SPC04 neural progenitor cells into neurons, diminished neurite outgrowth, and downregulated related genes, suggesting a MYT1L requirement for neural development; likewise, MYT1L knockdown during zebrafish neurogenesis altered hypothalamic development (Kepa et al. (2017); Blanchet et al. (2017)).

MYT1L is widely expressed in the developing brain in humans and mice, including cortical excitatory and inhibitory neurons. MYT1L expression is highest in post-mitotic neurons during fetal development but continues throughout later life (Kepa et al. (2017); Chen et al. (2021); Kim et al. (2022)), supporting roles in both brain development and function. Pathogenic mutations throughout the gene were reported, including heterozygous copy number variants, with gene deletion and nonsense, frameshift, and missense mutations resulting in IDDs, while duplications are associated with schizophrenia Pathogenic *MYT1L* variants contribute to a wide range of IDDs, including ASD, ID, obesity, behavioral disorders, facial dysmorphism, depressive disorder, and epilepsy (Blanchet et al. (2017); De Rocker et al. (2015); Coursimault et al. (2022); Windheuser et al. (2020); Mansfield et al. (2020)). Several studies in mouse models recently characterized consequences of MYT1L loss of function (LoF) mutation or deletion (Chen et al. (2021); Kim et al. (2022); Wöhr et al. (2022); Weigel et al. (2023)). These recapitulated some common human clinical phenotypes, including hyperactivity, impaired social interaction, repetitive activities, anxiety, and obesity. Two models exhibited precocious differentiation, disrupted neuronal maturation, and decreased synaptic gene expression (Chen et al. (2021); Weigel et al. (2023)), while other heterozygous LoF models did not exhibit precocious differentiation, but instead exhibited decreased expression of synapse-related genes (Kim et al. (2022)) or unaltered neurogenesis (Wöhr et al. (2022)). However, the consequences of pathogenic *MYT1L* mutation have not been characterized in human models.

Here, we generated and characterized the first human pluripotent stem cell models of *MYT1L* pathogenic mutation and employed a wide range of cellular, molecular, and functional assays to understand the etiology of IDDs stemming from these mutations. In the cortex, abnormal GABAergic inhibitory interneuron (cIN) and/or glutamatergic excitatory neuron (cExN) development contributes substantially to neurodevelopmental disorders (Gao and Penzes (2015); Zikopoulos and Barbas (2013)). Therefore, we used experimental paradigms for deriving cINs and cExNs from human pluripotent stem cell (hPSC) models to characterize the consequences of pathogenic *MYT1L* mutation, related these findings to its genome-wide occupancy, and compared these with the consequences of MYT1L deficiency. This work defined core aspects of altered neurodevelopment and function resulting from pathogenic *MYT1L* mutation, which constitute likely contributors to patient IDD phenotypes.

## RESULTS

### Generation and characterization of isogenic MYT1L variant hPSC models

We focused on an index proband subject with pathogenic *MYT1L* mutation and clinical characteristics of IDDs, including developmental delay, speech delay, moderate autism spectrum disorder (ASD) and intellectual disability (ID), lack of eye contact, repetitive behavior, attention deficit hyperactivity disorder (ADHD), depression, anxiety, and obesity (Table S1). Brain MRI findings were normal and there was no seizure history or facial dysmorphism (Table S1). To characterize the consequences of this variant, we derived pluripotent stem cell models from this proband (PD hPSC models) as described (Methods). We corrected the variant by genome engineering (PDC models) to generate isogenic pairs of models with and without the variant. We also derived isogenic models with and without this variant, by variant knock-in (VKI models) into wild-type hPSCs (WT). The subject carries a heterozygous single nucleotide duplication in the portion of the gene encoding the MYT1L domain, converting serine 707 to glutamine (S707Q) and predicted to cause a frameshift and premature stop codon (Fig. S1A). We validated the presence of the nucleotide duplication in the PD and VKI models (PD, VKI clonal lines 1/2) and its absence in isogenic controls (PDC, WT) (Fig. S1B). All models had a normal karyotype (Fig. S1C), had normal hPSC colony morphology and expressed markers of pluripotency (Fig. S1D).

To characterize consequences of pathogenic *MYT1L* mutation, all models were specified as cortical interneuron neural progenitor cells (cINPC, day (D) 15), differentiated into cortical interneurons (cINs, D30), and then further matured (m-cIN, D60), as described (Methods) (Meganathan et al. (2017); Meganathan et al. (2023)). All models were efficiently specified as medial ganglionic eminence-like neural progenitor cells (MGE-like cINPCs) with 87-92% expressing the MGE lineage marker NKX2-1 and NPC marker SOX1, as expected (Fig. S2). Thus, both variant and control models exhibited expected properties of pluripotent hPSCs and could be efficiently specified as MGE-like NPCs.

### The *MYT1L* S707Q variant results in haploinsufficiency

Abnormal GABAergic inhibitory interneuron (cIN) development can cause imbalanced neuronal function, contributing substantially to neurodevelopmental disorders (Gao and Penzes (2015); Zikopoulos and Barbas (2013)). As a recent study characterized the consequences of MYT1L heterozygous knockout in human cortical excitatory neuron-like cells generated by NGN2-mediated hPSC reprogramming (Weigel et al. (2023)), we assessed changes in MYT1L expression during human brain development and hPSC neuronal differentiation. During brain development, MYT1L expression increases from 8-12 post-conception weeks in the ventral and dorsal prefrontal cortex and persists throughout the lifespan (Fig. S3A; www.brainspan.org). During hPSC differentiation, MYT1L was not expressed in hPSCs (D0) or excitatory or inhibitory neural progenitors, but was expressed in newly differentiated cINs (D35) and excitatory neurons (cExNs (D40)) and was most highly expressed in matured neurons (m-cINs, m-cExN D60; Fig. S3B)(Meganathan et al. (2017); Chapman et al. (2024); Sanders et al. (2022)). During differentiation in neurosphere culture (D21), cINPC neurosphere size was similar in *MYT1L* variant and isogenic control models (Fig. S3C-C’). Relative to paired controls (PDC/WT), PD and VKI cINs exhibited significantly decreased MYT1L expression (Fig. 1A) and 30-47% protein level reduction (Figs. 1B-B’; Fig. S4A). Furthermore, a truncated protein product was not detected (predicted size 83.9 kDa), suggesting the mutant transcript underwent non-sense mediated decay as expected, and these models reflect MYT1L haploinsufficiency. Since MYT1L was highly expressed in both cINs and cExNs, we assessed whether the mutation affected NGN2-mediated hPSC reprogramming to cExN-like neurons and found that expression of both MYT1L and the pan-neuronal markers MAP2 and β-III TUBULIN was significantly reduced in the variant models (Fig. S4B-D), indicating that this variant affected neuronal gene expression in cExNs derived by NGN2-mediated hPSC reprogramming. However, as the consequences on cIN development remained entirely uncharacterized, we focused most subsequent analysis on understanding how *MYT1L* mutation affects this developmental program.

**Figure 1.**
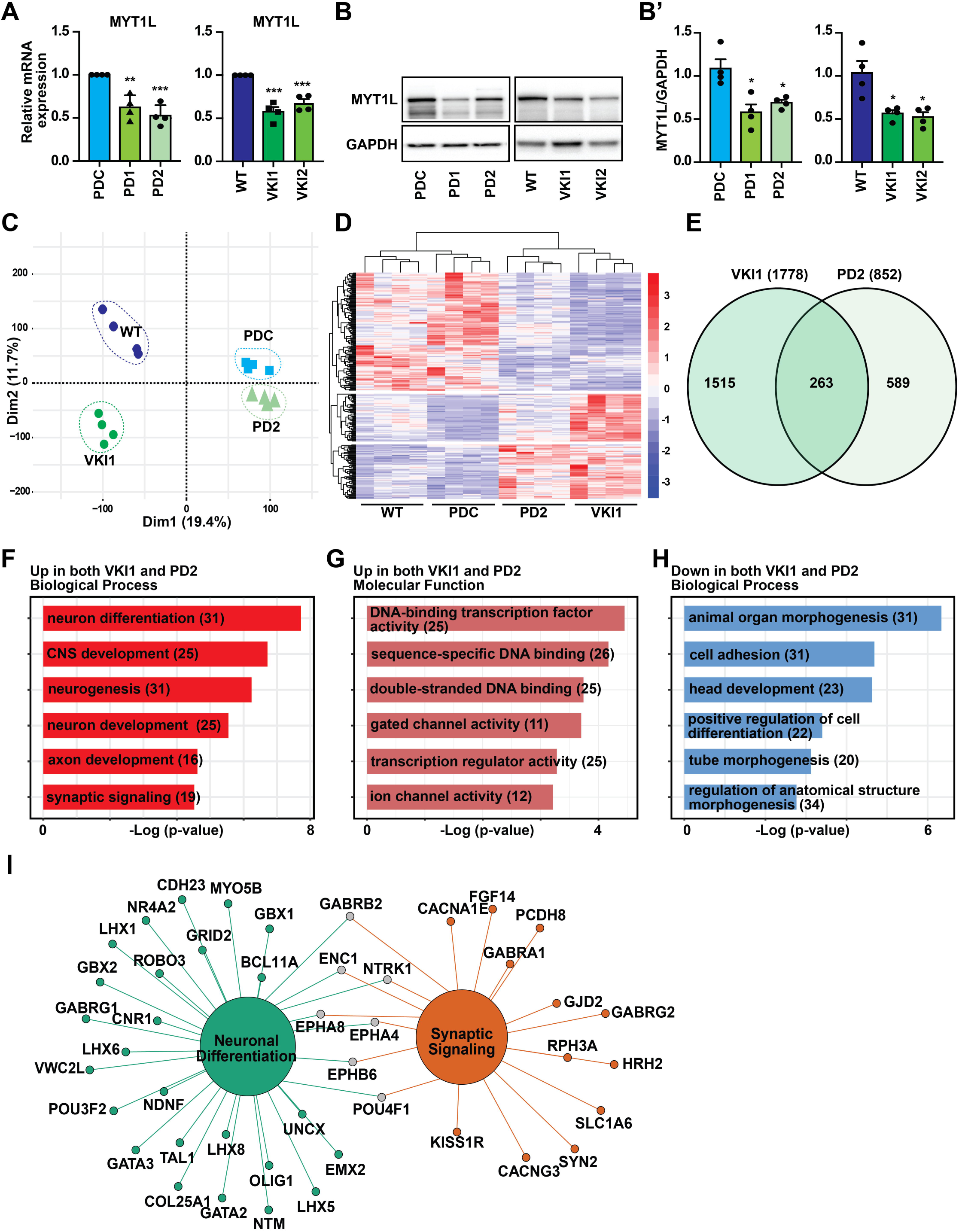
Transcriptomic analysis of hPSC-derived *MYT1L* S707QfsX56 variant cortical interneurons. **(A)** MYT1L expression was defined by RT-qPCR in cINs from the models shown. (**B**) MYT1L protein levels were defined by western blotting cIN protein lysates using a MYT1L-specific antibody (GAPDH as loading control). **B’** MYT1L protein levels, normalized to loading control. All four replicates are in Fig. S4A. (**C**) PCA plots visualize RNA-seq data for each model. (**D**) Expression levels of differentially expressed genes (DEGs) across 4 replicates each of cINs from these models. red=higher and blue=lower expression. (**E**) Venn diagram shows common significant DEGs for both VKI1-WT and PD2-PDC comparisons. (**F-H**) Gene ontology (GO) analysis of enriched (F) biological processes and (G) molecular functions for upregulated DEGs, and enriched (H) biological processes for downregulated DEGs common to the PD2 and VKI1 models. Number of enriched genes is in parentheses. (**I**) Network of upregulated DEGs under the neuronal differentiation and synaptic signaling GO terms in F. Green nodes, neuronal differentiation, orange nodes, synaptic signaling, and gray nodes, common genes. In all figures and panels, significance was calculated from 3-4 biological replicate experiments. Statistics were defined by two-tailed unpaired t-test, and quantitative data are shown as mean +/- SEM (Methods), with \**P* < 0.05, \*\**P* < 0.01, and \*\*\**P* < 0.001; for all figures and panels, actual values are in Table S9.

### Effects of the *MYT1L* variant on the human cortical interneuron transcriptome

To investigate the consequences of pathogenic *MYT1L* variation on the transcriptome, we performed RNA sequencing analysis on cINs derived from both variant and control models. Samples were generated for all four models and reads clustered by Principal Component Analysis (PCA)(Fig. 1C, Figs. S5A-B). Significantly differentially expressed genes (DEGs) were defined by data comparisons for each model-control pair (Methods)(Datasets S1A-B), with 852 and 1,778 DEGs defined for the PD2-PDC or VKI1-WT comparisons, respectively (Datasets S1C-D). These included 455 up- and 397 down-regulated DEGs (PD2-PDC) and 459 up- and 1,319 down-regulated DEGs (VKI1-WT)(Figs. S5C-D; Datasets S2A-B). 263 DEGs (122 up- and 141 down-regulated) were commonly dysregulated in both variant models versus their isogenic controls (Figs. 1D-E; Dataset S1E; Dataset S2C). Gene ontology (GO) enrichment analysis of up-regulated DEGs in either the PD2, VKI1, or both datasets revealed biological process (BP) terms related to neuronal differentiation and synaptic signaling (Fig. 1F, S5E-F; Datasets S3A-C) and up-regulated Molecular Function (MF) pathways related to transcription and neuronal function (Fig. 1G, S5E-F; Datasets S3A-C). By contrast, down-regulated genes enriched for general biological processes related to early development (Fig. 1H; Datasets S3D-F), and general molecular function GO terms (e.g. calcium ion or growth factor binding; Datasets S3D-F).

As many upregulated DEGs related to neuronal differentiation and synaptic signaling, we performed network analysis on genes in the GO categories from Fig. 1F (Fig. 1I), defining related gene networks (Fig. S5G-H). These data indicate that the *MYT1L* variant models upregulate genes related to axon development and synaptic signaling versus their isogenic controls. We confirmed up-regulated expression in the PD2 and VKI1 models of suites of ‘neuron differentiation’- and ‘synaptic signaling’-related genes (Figs. 2A-B) and validated a subset by RT-qPCR (Figs. 2C-D).

**Figure 2.**
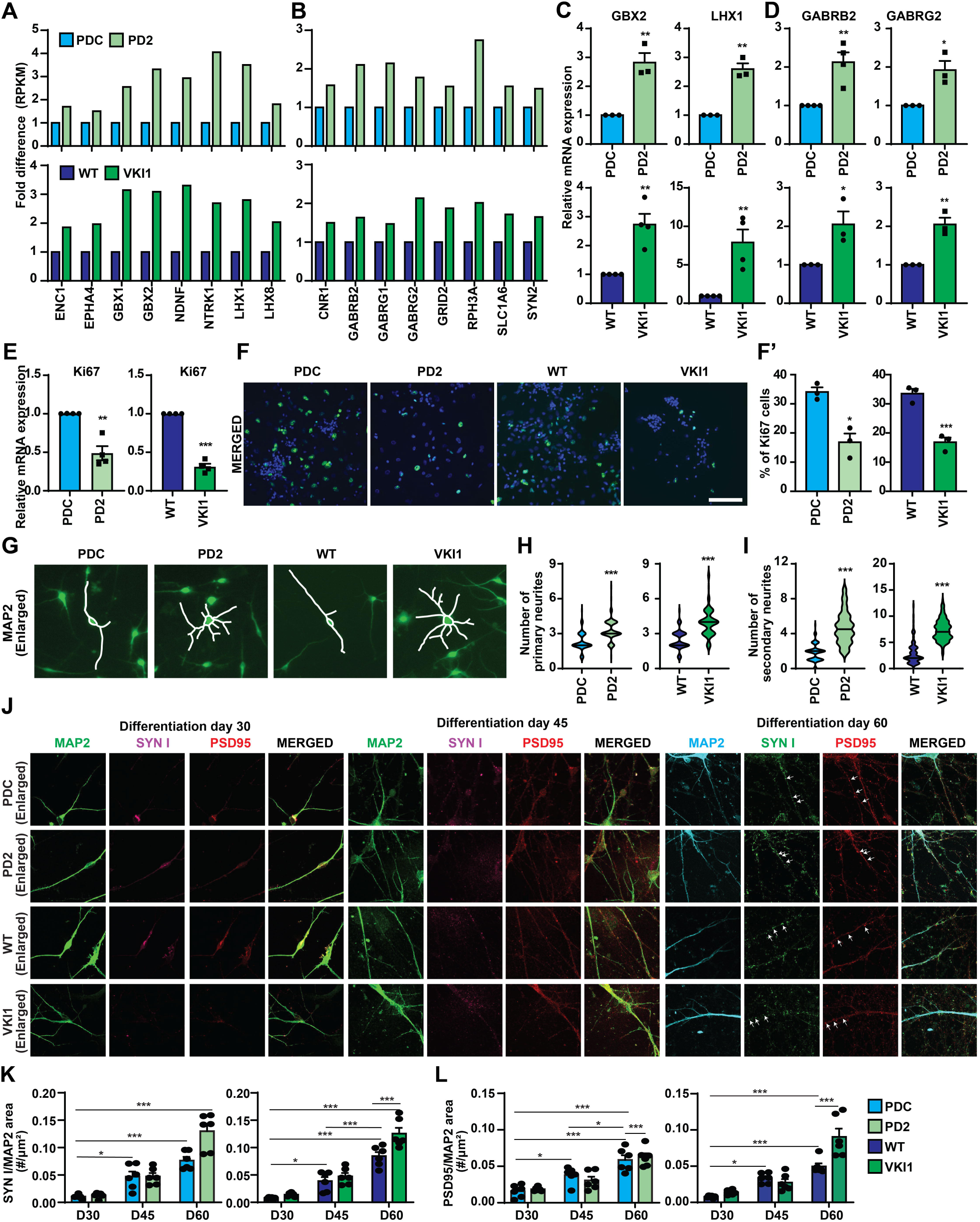
*MYT1L* variant alters the neuronal differentiation and synaptic signaling gene expression and neuronal morphology. (**A-B**) DEGs involved in (A) neuronal differentiation and (B) synaptic signaling in the PDC-PD2 models (top) and WT-VKI1 models (bottom). Fold differences in expression of significant DEGs are represented as average reads per kilobase of transcript per million mapped reads (RPKM). (**C-D**) Relative expression of the genes shown was defined by RT-qPCR in cINs. (**E**) Ki67 expression, detected by RT-qPCR. (**F**) Ki67 immunostained cINs (scale bar=100 µM). **F’**: percentage of Ki67-positive cells (extended images in Fig. S6). (**G**) Enlarged images from Fig. S6J show MAP2 stained cINs, with tracing of cell soma, primary, and secondary neurites. (**H**) Primary and (**I**) secondary neurites were counted in cINs across 3 biological replicates and 40 cINs per replicate for each model. (**J**) Enlarged images from Fig. S6K of MAP2, SYN I, and PSD95 immunostained cINs. Arrows indicate SYN I and PSD95 puncta. **(K-L)** Quantification of (K) SYN I and (L) PSD95 puncta per MAP2 stained area, quantified with MATLAB Intellicount software. Significance was calculated by two-tailed unpaired t-test or 2way ANOVA with Tukey’s multiple comparison, as mean with +/- SEM. \**P* < 0.05, \*\**P* < 0.01, and \*\*\**P* < 0.001.

### Enhanced differentiation and altered morphology in *MYT1L* pathogenic variant models

As the increased neuronal and synaptic gene expression in variant models suggested altered cIN differentiation, we examined the proliferation marker Ki67 in cultures of cINs differentiated in the presence of the Notch pathway inhibitor DAPT (Methods). Both variant models exhibited significantly reduced Ki67 mRNA expression (Fig. 2E) and a nearly 50% decrease in the Ki67-immunopositive cell fraction (Fig. 2F-F’; Fig. S6). As >30% of the day 30 control cells were still Ki67-positive when using this differentiation protocol, we repeated this experiment with several protocol variants, performing differentiation with the addition of no small molecules, DAPT, PD0332991 (a CDK4/6 inhibitor), or both (PD+DAPT). As expected, a similar Ki67-positive cell fraction was observed as above with no small molecule or DAPT treatment and this was significantly reduced in *MYT1L* variant models versus controls (Fig. S6A-E). Differentiation in the presence of PD or PD+DAPT instead yielded a very low fraction of Ki67-positive cells (∼2%) in the control models, such that further reduction in the *MYT1L* variant models could not be assessed (Fig. S6A-E). Together, these findings indicate that *MYT1L* mutation reduces the fraction of proliferating cells, coincident with increased expression of neuronal differentiation and synapse-related genes.

As variant cINs exhibited enhanced neuronal and synaptic gene expression, we also tested whether this altered cIN morphology by re-seeding cINs at low density on D25, continuing differentiation through D30, and quantifying primary neurites associated with each cell soma (Fig. S6F). Most control cINs exhibited bipolar morphology, with two primary neurites, while variant neurons had increased numbers of primary neurites and a multipolar morphology (Fig. S6F-G). Secondary neurites originating from primary neurites also increased in variant cINs (Fig. S6F, H), while PD2 cell soma were also significantly larger than controls (Fig. S6I). To confirm that these neurites originated from neurons, we quantified them in MAP2 immunostained cINs on D30 (Fig. 2G-I; Fig. S6J). Therefore, *MYT1L* variant models exhibited altered differentiation and morphology, with production of more primary and secondary neurites. We also examined whether this increased morphological complexity was accompanied by altered production of synaptic puncta expressing the pre-synaptic gene SYNAPSIN I (SYN I) and post-synaptic gene PSD95, performing immunohistochemistry at D30, 45, and 60. Puncta expressing these markers were detected at D45-60 and were significantly more abundant in variant neurons (Fig. 2J-L, S6K), suggesting that the increased synaptic gene expression may also enhance production of puncta expressing pre- and post-synaptic markers.

### *MYT1L* deficiency in the variant models diminishes cortical interneuron gene expression

We examined how the *MYT1L* variant could affect proliferation and differentiation-related marker expression by initially assessing endogenous expression of Ki67 and the cINPC marker NESTIN, neuronal marker MAP2, and cIN marker somatostatin (SST) during differentiation into m-cINs (Fig. S7A)(Meganathan et al. (2017); Chapman et al. (2024)). NESTIN and Ki67 expression peaked in NPCs, gradually declined during differentiation (D30), and was almost undetectable in m-cINs (D60), while MAP2 and SST expression peaked at D30 and D60 respectively. We then used PD0332991+DAPT treatment (rather than DAPT alone as in Fig. S6A-E) to more efficiently differentiate the variant and control lines. With this treatment, only a small percentage of control cells (∼2%) expressed NESTIN or Ki67 at D45-60 (Fig. S7B-E), and there was a trending but insignificant further reduction of the proliferating cell fraction in the variant models. In these experiments, the MAP2-immunopositive fraction was also significantly reduced in the *MYT1L* variant models at D45-60 (Fig. S7E). *MYT1L* variant cINs also exhibited significantly reduced expression of the pro-neuronal marker ASCL1, cIN migration marker DLX2, cIN markers calbindin 1 (CALB1), CALB2 (calretinin), and SST, and pan-GABAergic neuron markers GAD1 and GAD2 (Fig. 3B-C). We further validated these findings by immunostaining, finding that the CALB1-immunopositive cell fraction was also significantly reduced in the variant models (Fig. 3D-D’), indicating that cortical interneuron and GABAergic neuron marker expression was impaired in the *MYT1L* variant models during differentiation.

**Figure 3.**
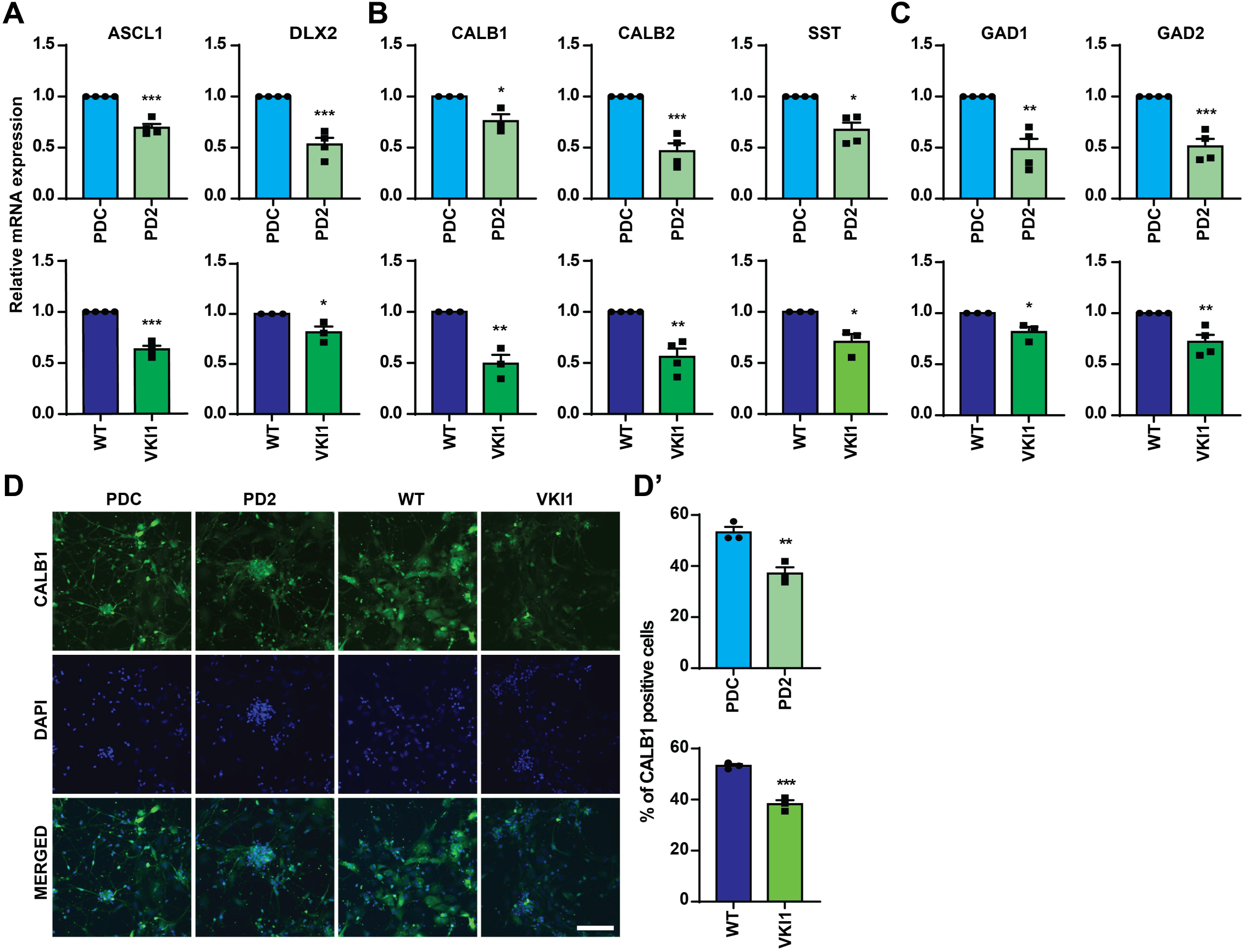
Expression of proneuronal and cortical interneuron maturation markers is diminished in cINs carrying the *MYT1L* variant. **(A-C)** Expression of the (A) proneuronal and (B-C) mature cortical interneuron markers shown, assessed by RT-qPCR in cINs. (**D**) Representative images of cINs immunostained for CALB1, and DAPI counterstained (scale bar=100 µM). **D’** Percentage of CALB1-positive cells, relative to DAPI-positive nuclei. Significance was calculated by two-tailed unpaired t-test as mean +/- SEM. \**P* < 0.05, \*\**P* < 0.01, and \*\*\**P* < 0.001.

### MYT1L CRISPRi knockdown compromises neurite outgrowth and expression of mature neuronal and interneuron marker gene expression

To further investigate MYT1L requirements during cortical interneuron development, we generated CRISPR inhibition models. MYT1L promoter-specific guide RNAs (g1/2) were transduced into WT cINPCs stably expressing a nuclease deactivated Cas9-KRAB repressor domain fusion (dCas9-KRAB). Lines were differentiated into D30 cINs to assess consequences of knockdown (KD). Both gRNAs reduced MYT1L mRNA expression levels more substantially than our variant models above, by 68% or 72% respectively, versus the dCas9-KRAB control (Fig. 4A). We plated cINPC spheres during differentiation and found that both KD models exhibited significantly reduced neurite outgrowth versus controls (Fig. 4B-B’). Day 30 cINs from both KD models exhibited significantly reduced expression of the pan-neuronal markers MAP2 and β-III TUBULIN (Fig. 4C). Likewise, both models exhibited significantly reduced expression of cIN differentiation markers DLX2 and DCX, and CALB1, CALB2, SST, GAD1, and GAD2 (Fig. 4D). We further found that the cell fractions immunopositive for MYT1L, β-III TUBULIN, MAP2, and SST were significantly reduced in the KD for all four markers (Fig. 4E-E’). These data indicate that MYT1L deficiency impairs the expression of general neuronal, cIN, and GABAergic neuron markers, which may compromise cIN maturation or identity.

**Figure 4.**
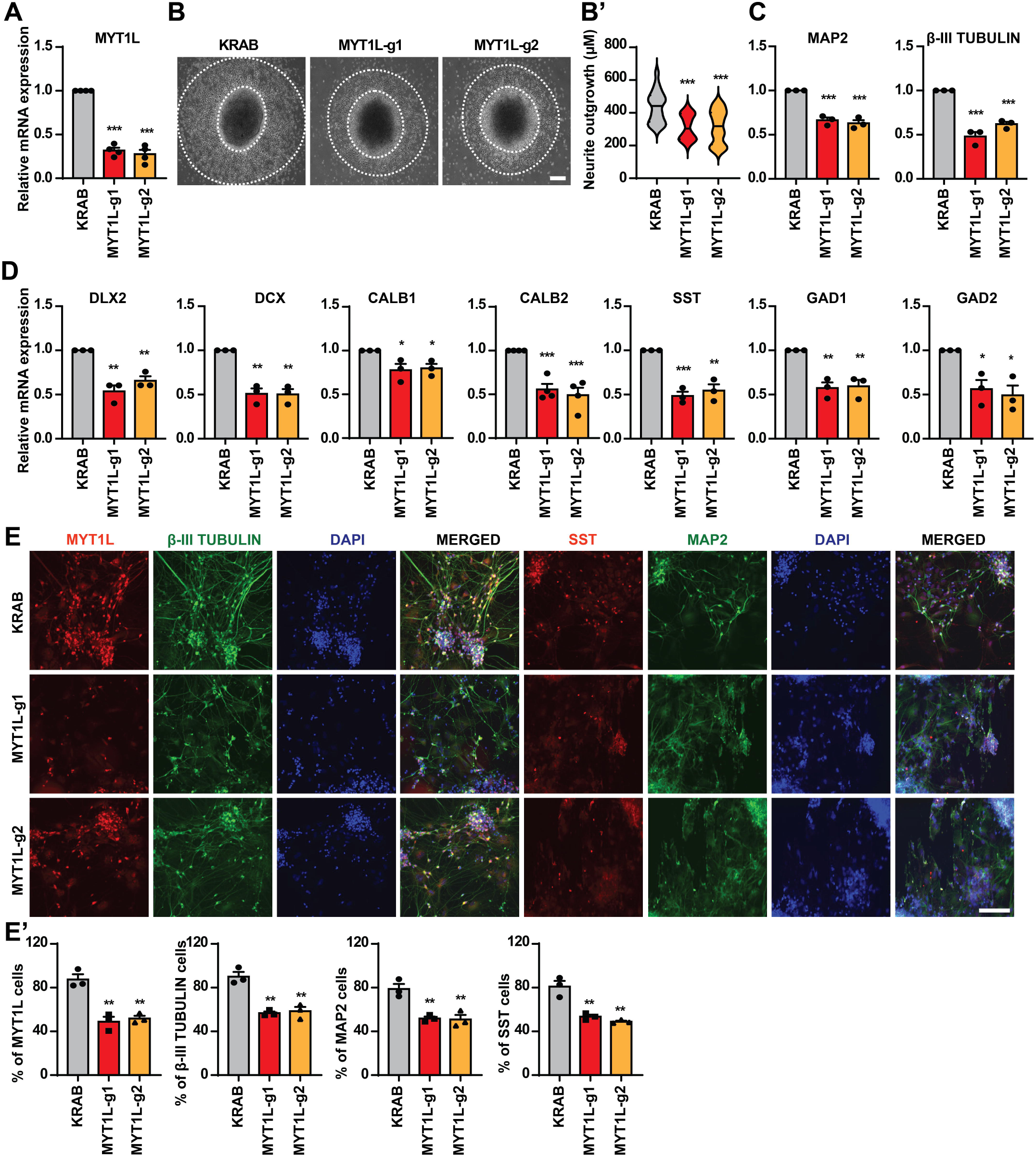
MYT1L knockdown compromises neurite outgrowth and neuronal differentiation and maturation marker expression. **(A)** MYT1L expression defined by RT-qPCR in cINs expressing dCas9-KRAB (KRAB control) +/- gRNAs. (**B**) Representative images of neurite outgrowth in plated D22 spheres. Scale bar=200 µM. Neurite outgrowth is highlighted with dotted lines. B’: quantification, with 14 spheres assayed per replicate. (**C-D**) Expression of the markers shown, defined by RT-qPCR for the same samples. (**E**) Representative 20X images of cINs immunostained for MYT1L and β-III TUBULIN or SST and MAP2, and DAPI counterstained. Scale bar=100 µM. Data shown was obtained from three biological replicate experiments with significance calculated by two-tailed unpaired t-test and quantitative data shown as mean+/-SEM. *P* values: \**P* < 0.05, \*\**P* < 0.01, and \*\*\**P* < 0.001.

### Matured cortical interneurons with the *MYT1L* variant diminished expression of pan-neuronal and cortical interneuron marker genes

As we observed reduced cIN marker expression in both the *MYT1L* variant and CRISPRi KD models at D30, we also assessed whether *MYT1L* deficiency and mutation also affected cIN maturation to D60. This was warranted because MYT1L expression levels are substantially higher in D60 m-cINs, versus newly differentiated D30 cINs (Fig. S3B). MYT1L expression levels were reduced by 50% in m-cINs derived from both variant models, relative to paired controls (Fig. 5A). In addition, m-cINs exhibited significantly reduced expression of the neuronal markers MAP2, β-III TUBULIN (Fig. 5B), cIN markers CALB1, CALB2, and SST, and GABAergic neuron markers GAD1 and GAD2 (Fig. 5C-D).

**Figure 5.**
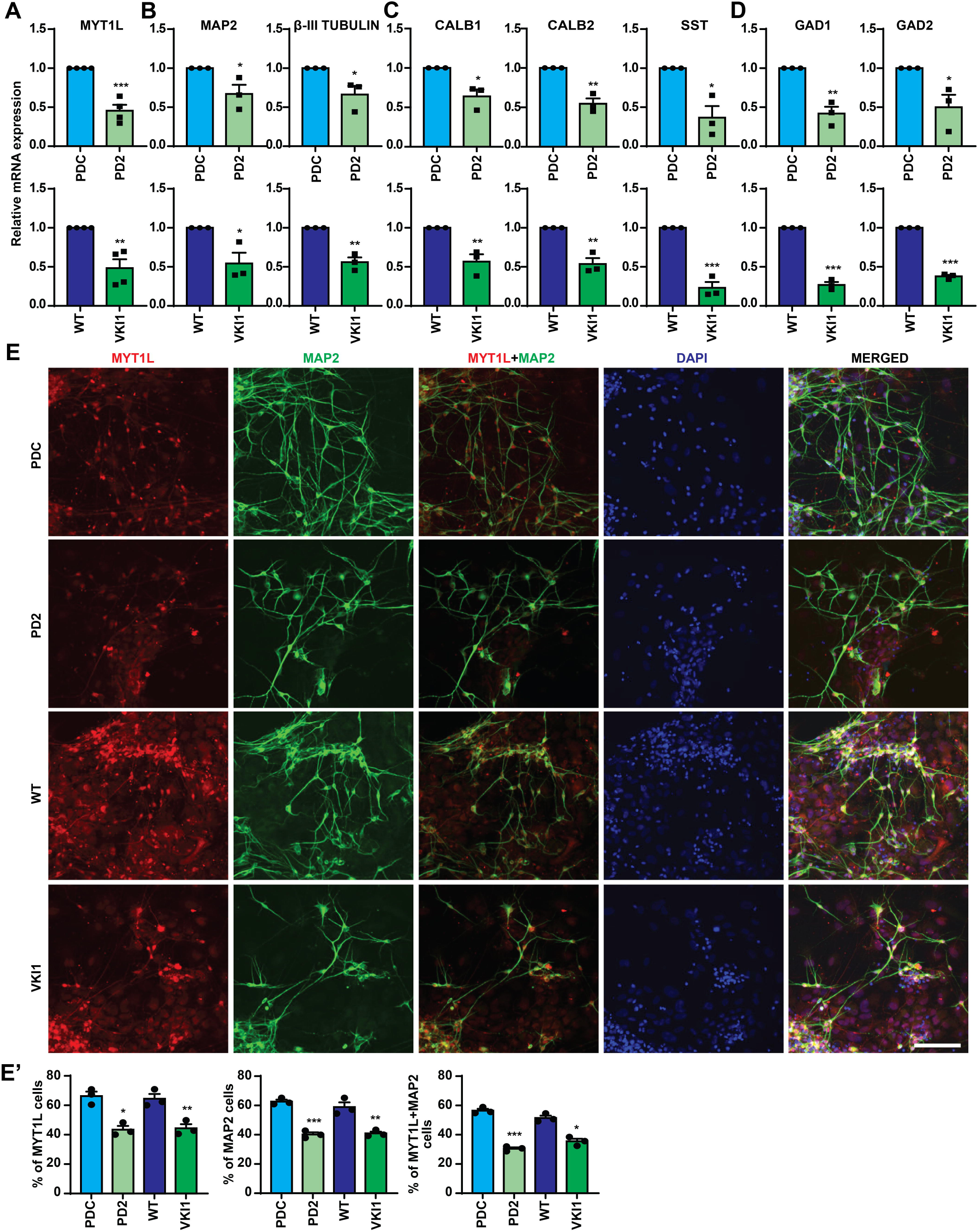
Expression of markers of neuronal and cortical interneuron maturation and identity are diminished in *MYT1L* variant m-cINs. (**A-D**) Expression of genes shown was defined by RT-qPCR in m-cINs. (**E**) Representative images of m-cIN immunostained for MYT1L and MAP2, and DAPI counterstained (scale bar=100 µM). **E’**: percentage of MYT1L, MAP2, and MYT1L+MAP2 (double) positive cells, with significance calculated by two-tailed unpaired t-test as mean +/-SEM. \**P* < 0.05, \*\**P* < 0.01, and \*\*\**P* < 0.001.

To assess whether these findings could reflect altered cell fractions of cells expressing these markers, we conducted immunocytochemistry for MYT1L and MAP2 in D60 m-cINs. As expected, the fraction of cells immunopositive for MYT1L at the threshold set in ImageJ (see Methods) was significantly reduced in the variant models, and the MAP2 positive and MYT1L/MAP2 double immunopositive cell fractions were likewise significantly reduced in variant versus control models (Fig. 5E-E’). Together, these data indicate that neuronal and cortical interneuron gene expression is diminished during interneuron differentiation and maturation in models with a *MYT1L* pathogenic variant.

### Electrophysiological function and channel gene expression and activity is disrupted in *MYT1L* variant interneurons

To assess whether the *MYT1L* variant affected cIN function, we next conducted electrophysiology, recording cells differentiated from the VKI1 and isogenic control line (WT)(Fig. 6). D30 neurons were plated on cortical rat astrocytes and recordings were performed from D60 (Fig. 6; Table S8). Cells of both genotypes expressed tetrodotoxin (TTX) sensitive inward sodium (Na) currents and 4-Aminopyridine (4-AP) sensitive transient and tetraethylammonium (TEA) sensitive, sustained outward potassium (K) currents (Fig. 6A). However, the VKI1 m-cINs at 4 weeks after plating exhibited significantly smaller voltage-gated currents than isogenic controls, including diminished Na current and both transient and sustained K currents (Fig. 6A-F; Table S8). In addition, VKI1 cells displayed a slightly more negative potential for steady-state current reversal (Fig. 6G; Table S8).

**Figure 6.**
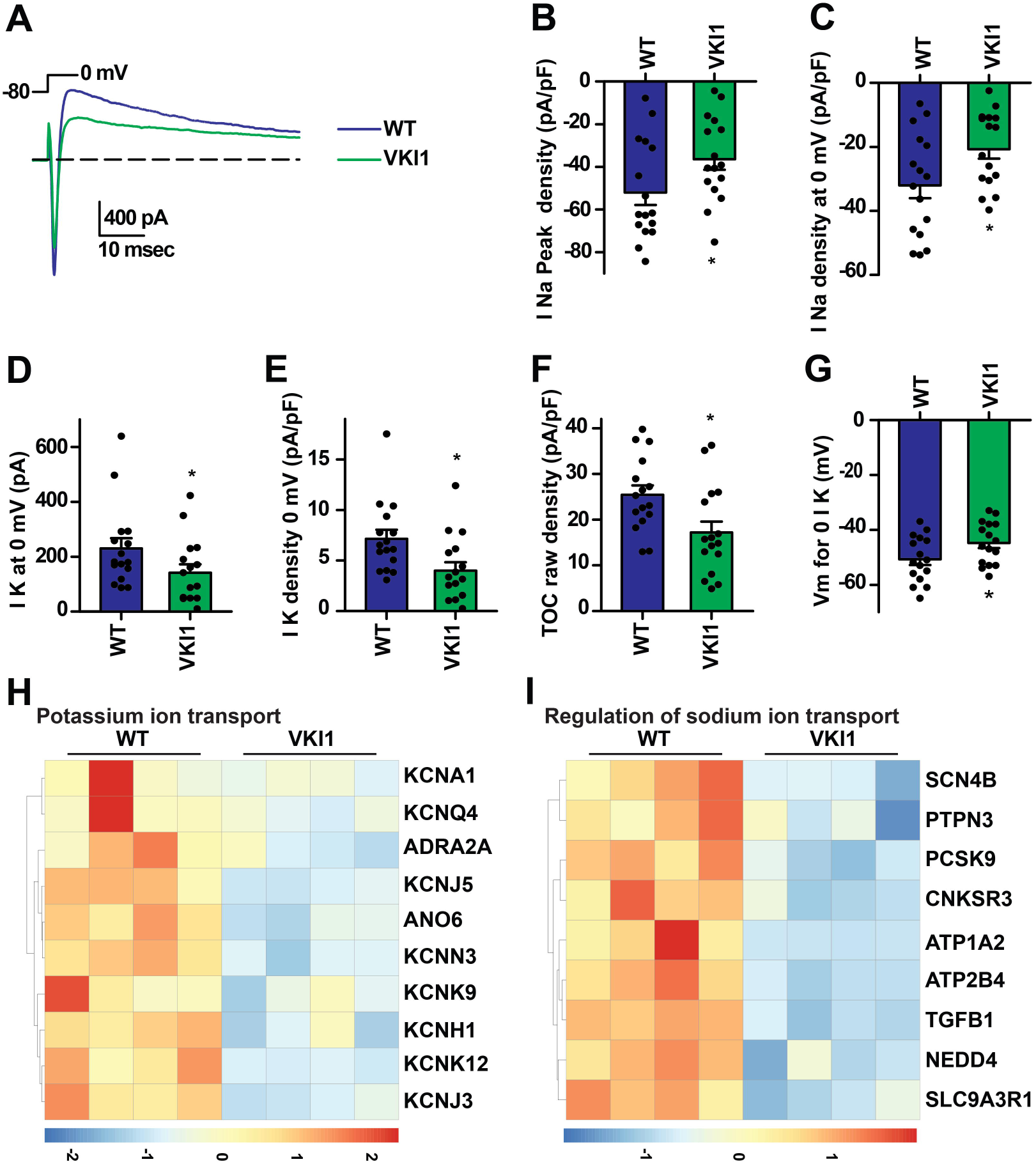
*MYT1L* variant m-cINs exhibit altered voltage-gated currents. **(A)** Inward and outward currents evoked by voltage step from -80-0 mV in neurons differentiated from WT (blue) and VKI1 (green). (**B-E**) VKI1 cells had (B) reduced tetrodotoxin (TTX) sensitive inward sodium (Na) current, (D) reduced tetraethylammonium (TEA) sensitive steady-state outward potassium (K) current, and (C, E) reduced normalized current densities. In addition, VKI1 cells displayed (**A, F**) less 4-aminopyridine (4-AP; a potassium channel antagonist) sensitive transient outward current and (**G**) less negative reversal potential for steady-state current. (**H-I**) DEGs in VKI1-WT comparisons under the GO terms (H) potassium ion transport and (I) regulation of sodium ion transport. Significance was calculated by two-tailed unpaired t-test as mean +/-SEM. \**P* < 0.05. with actual values in Table S8.

To assess whether these functional changes in sodium and potassium currents were related to altered cIN maturation or gene expression, we assessed differentially expressed genes involved in potassium ion transport and regulation of sodium ion transport, based upon genes under these GO terms in our transcriptomic datasets (Dataset S3F). Most of these genes were down-regulated in the VKI model cINs versus the isogenic control (WT) (Fig. 6H-I; Dataset S2B), suggesting that reduced ion transport-related gene expression could contribute to the reduced sodium and potassium currents seen in our electrophysiological analysis.

### Genome-wide MYT1L occupancy and integration with differentially expressed genes in *MYT1L* variant model interneurons

To understand mechanisms by which MYT1L could regulate cIN gene expression, we also determined its genome-wide occupancy by performing Cleavage Under Targets & Release Using Nuclease (CUT&RUN) in D30 cINs (Skene and Henikoff (2017), identifying reproducible peaks present in all four MYT1L CUT&RUN biological replicate experiments and not the IgG control (see Methods; Fig. 7A; Dataset 4A). We annotated peaks to the nearest transcription start site and found that the majority mapped either to promoters or distal intergenic or intronic locations, which might act as cis-regulatory elements, with few located in other genomic locations (3’ UTR, 5’ UTR, exonic; Fig. 7B, Dataset 4A). We assessed chromatin state underlying these MYT1L bound peaks by ChromHMM partitioning the genome from D0-60 and clustering for four histone modifications (H3K4me3, H3K27me3, H3K27ac, H3K9me3), as described previously (Chapman et al. (2024))(Fig. 7C; Dataset 4B). Most MYT1L bound peaks (60%) co-enriched with H3K27ac and H3K4me3, histone modifications associated with active chromatin (Fig. 7C; Dataset 4B; clusters 2-4), while a minority of targets associated with bivalent chromatin (cluster 5, marked by H3K27me3 and H3K4me3; 11%) or repressive chromatin marked by H3K27me3 or H3K9me3 (clusters 6-7; 17%)(Fig. 7C-D; Dataset 4B). We mapped each MYT1L bound peak to the nearest transcription start site and analyzed the associated genes (Dataset S4A). Top pathway enrichment GO terms were related to transcription (Fig. 7E; Dataset 4C), while associated biological processes and molecular functions also related to transcription, indicating that putative MYT1L direct targets enrich for genes with transcription regulator activity (Fig. 7E; Dataset S4C). We also observed enriched GO terms related to neuronal and synaptic function (Fig. 7F-G; Dataset S4C) suggesting that MYT1L acts by directly regulating the expression of both neuronal and synaptic genes and transcription factors that may also control these processes.

**Figure 7.**
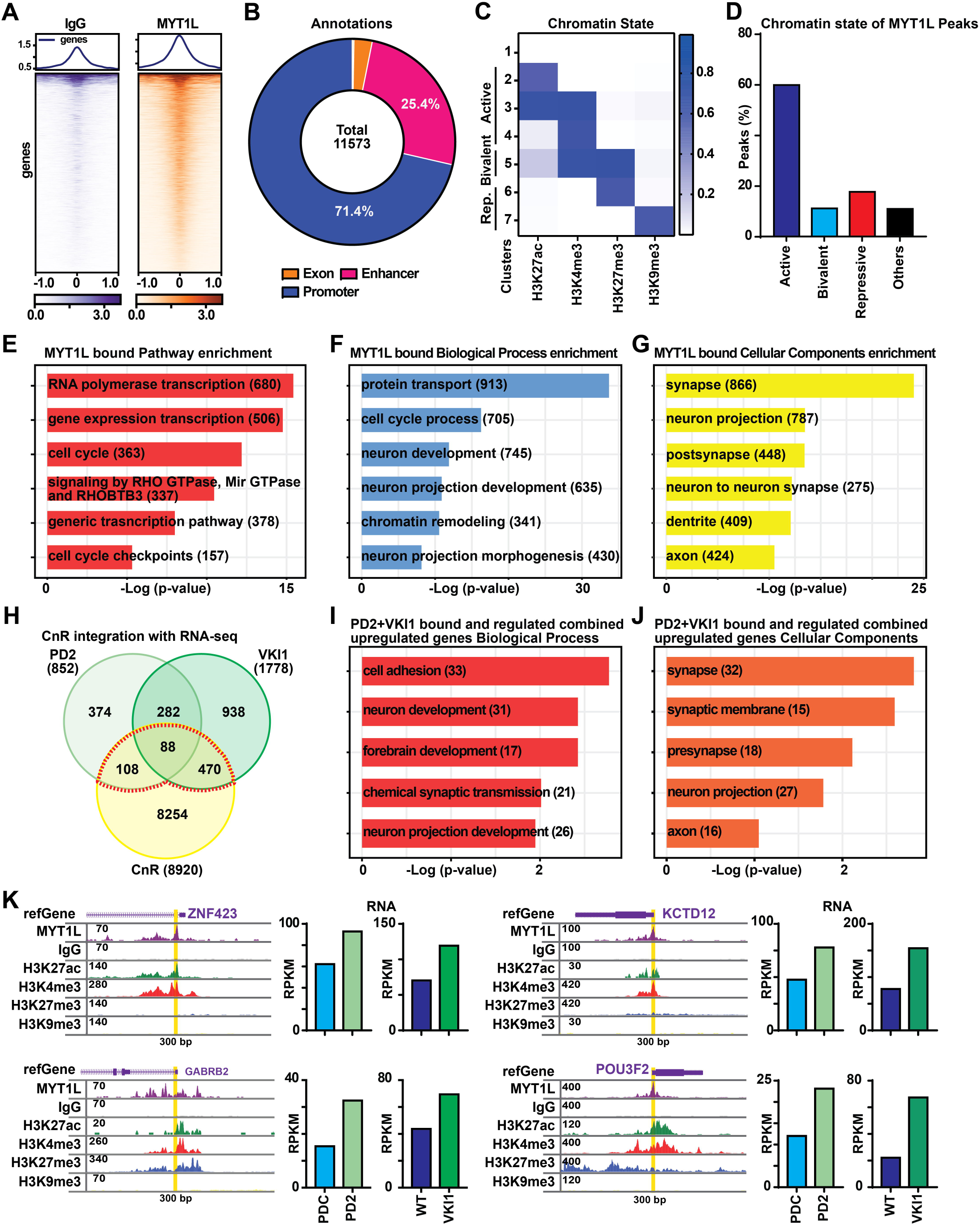
Genome-wide MYT1L occupancy and integration with differential gene expression in variant cortical interneurons. **(A)** Deep tools visualizations of MYT1L bound peaks and IgG signal at the same locations (n=4 (MYT1L), n=3 (IgG) replicates).). **(B)** MYT1L peak annotations and **(C)** Chromatin states, defined by ChromHMM partitioning, based upon aggregate enrichment for four histone modifications from D0-D60; heatmap shows emission probabilities for the 7 clusters. **(D)** MYT1L bound peaks associated with each chromatin state. **(E-G)** GO enrichment of MYT1L bound genes. **(H)** Overlap between MYT1L bound genes and significant DEGs. **(I-J)** GO analysis of genes associated with MYT1L bound peaks and significantly upregulated in variant cINs. **(K)** Browser tracks show MYT1L bound peaks and associated histone modification state for representative genes in cINs, with differential gene expression (RPKM values; right).

We next integrated these MYT1L bound genes with significant DEGs in the *MYT1L* variant versus control data, defining 169 PD2 and 475 VKI1 DEGs as both bound and regulated (Fig. 7H; Dataset S5A). In both comparisons, genes associated with MYT1L binding and upregulated in the variant versus control models enriched for GO terms related to neuron development and synaptic signaling (Datasets S5B-C). We also assessed MYT1L binding-associated genes that were down-regulated in variant models, but few genes met these criteria and significant GO terms were not obtained for this limited gene set (Datasets S5D-E). We then combined all MYT1L bound DEGs (for either comparison) and used this gene set (highlight, Fig. 7H) for GO term analysis. These combined bound and upregulated genes were enriched for neurodevelopmental GO terms, including ‘forebrain development’, ‘synapse’, and ‘neuron projection’ (Fig. 7I-J; Datasets S5F-G). We visualized several MYT1L bound peaks associated with DEGs as browser tracks, where MYT1L binding was associated with either active (ZNF423/KCTD12) or bivalent (GABRB2/POU3F2) chromatin states, while all four genes increased in expression in variant versus control cINs (Fig. 7H).

Finally, to define the most likely IDD contributory MYT1L direct targets, we assessed which MYT1L bound significant DEGs were high confidence ASD or epilepsy genes, as these are common comorbidities in individuals with *MYT1L* mutation (Simons Sfari; 1,076 genes; 07-20-2022 release; Dataset S6A) and EpilepsyGene (499 genes; 05-22-2023 download; Dataset S6B). In these analyses, both up- and down-regulated MYT1L direct target genes were enriched for high confidence ASD and/or epilepsy genes (Fig. S8A-B). Many of these have known roles in neurodevelopment, including *GABRB2*, *NR4A2*, *SLC4A10*, *CSMD3*, *RELN*, *CNR1*, *CNTN5*, *BCL11A*, and *NRP2*. These data indicate that MYT1L is required for transcriptional control of a network of downstream genes involved in neuronal maturation and function, with dysregulated expression of other IDD genes likely contributing to phenotypes in individuals who carry pathogenic *MYT1L* mutation. In summary, these data indicate that *MYT1L* pathogenic mutation affects interneuron differentiation, maturation, and function (Fig. S8C), effects that may relate to MYT1L-mediated direct regulation of IDD-relevant and other neuronal genes implicated in these processes.

## DISCUSSION

Pathogenic *MYT1L* mutations are a recently defined contributor to IDDs, with most patients having heterozygous *de novo* or, in rare cases, inherited predicted loss of function mutations associated with IDD phenotypes including ASD, ID, and epilepsy. However, alterations in human neurodevelopment and neuronal function stemming from pathogenic mutation had not been characterized. Therefore, here we generated the first human cellular models of pathogenic *MYT1L* mutation, which exhibited haploinsufficiency for MYT1L RNA and protein. Upon inducing differentiation of these variant models into cortical interneurons, neuronal and synapse-related gene expression increased, with a corresponding decrease in proliferating progenitors, mutant neurons exhibited more complex morphology, with increased primary and secondary neurites, and synaptic puncta expressing pre- and post-synaptic markers. However, during neuronal maturation, expression of markers of neuronal maturation and cortical interneuron identity was reduced, while electrophysiological analysis demonstrated reduced sodium and potassium channel activity. Together, our data indicates that pathogenic *MYT1L* mutation disrupts cortical interneuron differentiation, maturation, and function.

During human fetal development, MYT1L expression is initiated during the cell cycle exit accompanying neuronal differentiation, with levels peaking in the fetal and early postnatal brain but persisting throughout the lifespan (Kepa et al. (2017); www.brainspan.org), while Myt1l is widely expressed in different brain regions in the mouse, further supporting a role in brain development and later function (Chen et al. (2021); Kim et al. (2022)). Here, we developed multiple models carrying the mutation present in our index MYT1L patient (c.2117dupG; MYT1L S707fsX). This is analogous to the amino acid residue mutated in the first Myt1l mouse model published recently by Chen et al (2021), while additional Myt1l mouse models were also recently reported targeting exons 6 (Wöhr et al. (2022); Weigel et al. (2023)) or 9 (Kim et al. (2022)). Animals heterozygous for these deletions or mutations all exhibited reduced Myt1l mRNA and protein, modeling the haploinsufficiency typical of patients and seen in our hPSC models.

While differences in the analyses and time frames conducted preclude comprehensive comparisons between our human and the mouse models, both similarities and differences were observed. Distinct consequences on neurogenesis are reported in mouse models, with Chen et al. reporting that Myt1l homozygous loss of function in embryonic (E14.5) cortex caused upregulation of neuronal differentiation and immature neuronal marker genes and depletion of cells expressing stem cell and proliferative markers, congruent with findings in our hPSC models (Chen et al. (2021). However, in other studies, Myt1l deficiency increased expression of the neural stem cell marker Sox2, while cortices were thinner at birth, suggesting impaired neurogenesis (Weigel et al. (2023)) or no effects on neurogenesis at E18.5 were identified (Wöhr et al. (2022)), while these models displayed some shared behavioral phenotypes at later time points (Chen et al. (2021); Weigel et al. (2023)). While our findings are most consistent with Chen et al. (2021), the discrepant findings in mouse studies could relate to assessing heterozygous versus homozygous animals at different time points, or to differences between the mouse models.

Our heterozygous models of *MYT1L* mutation exhibited downregulated expression of markers of neuronal maturation and cortical interneuron identity, both at earlier (D30) and later (D60) time points after initiating neuronal specification and differentiation, indicating a major, requirement for normal MYT1L dose for neuronal maturation, maintenance of identity, and function. Work in two mouse models are generally congruent with these findings: neuron projection morphogenesis and potassium ion transport were downregulated in the adult prefrontal cortex of one mouse model, suggesting disruption of neuronal maturation and persistence of immature neuronal marker expression (Chen et al. (2021)). Likewise, another heterozygous knockout mouse model exhibited decreased striatal and hippocampal synaptic gene expression, indicating impaired neuronal maturation at adult but not juvenile (P21) stages, supporting late onset (Kim et al. (2022).

In prior work, Myt1l overexpression could promote neuronal conversion and maturation, while shRNA knockdown reduced neurite outgrowth and neuronal maturation (Vierbuchen et al. (2010); Mall et al. (2017); Kepa et al. (2017)). We likewise found that CRISPRi-mediated MYT1L deficiency during cIN differentiation hampered neurite outgrowth and reduced expression of pan-neuronal, cIN, and GABAergic neuron markers, indicating a MYT1L requirement for human cIN maturation. Unlike our CRISPRi deficiency models, which reduced MYT1L mRNA levels by 68-72%, pathogenic *MYT1L* haploinsufficiency in our variant models did not impair initial neurite outgrowth from plated neural progenitor spheres. However, in both models, cortical interneuron marker expression was diminished, suggesting a core MYT1L requirement for neuronal maturation and acquisition of mature cIN identity. The observation of impaired neuronal maturation both in our human and prior mouse models supports this as a core phenotype contributing to altered neuronal function.

While functional consequences of MYT1l mutation were largely uncharacterized in human models, one recent study assessed altered function in cortical excitatory-like neurons generated by NGN2 overexpression-mediated reprogramming (Weigel et al. (2023)). These neurons exhibited impaired neurogenesis, by contrast with our findings for interneurons and with some mouse models (e.g. Chen et al. (2021)). Reminiscent of our findings for cINs, these NGN2-induced excitatory neurons showed impaired synaptic gene expression, perhaps indicating impaired maturation, but after 6 weeks of culture exhibited hyperexcitability (Weigel et al. (2023), while our D60 cINs instead exhibited diminished neuronal activity and correspondingly reduced sodium and potassium channel expression and activity. These differences may relate to differences in neuronal type, derivation methods, or timing.

We intersected *MYT1L* mutation-related transcriptomic dysregulation in cINs with its genome-wide occupancy, demonstrating enrichment for genes involved in transcriptional regulation and synaptogenesis. Like similar results for mouse cortical neurons (Mall et al. (2017); Chen et al. (2023)), MYT1L binding in human cINs was predominantly associated with active histone modifications around gene promotors and putative enhancers in intergenic locations. MYT1L bound and down-regulated putative direct targets include synaptic genes that are also mutated to cause IDD and may contribute to impaired neuronal maturation and function seen in our *MYT1L* mutant models. For example, *BCL11A* mutation is associated with cognitive and motor delay and ASD (Bruce and Peter (2022)), and de novo or inherited mutations in the mitochondrial membrane protein GPD2 contribute to ID and ASD (Lewis et al. (2019); Barge-Schaapveld et al. (2013)). Dysregulation of these and other genes stemming from pathogenic MYT1L mutation may contribute to IDD phenotypes.

Here, we showed that *MYT1L* pathogenic mutation resulted in haploinsufficiency. While consistent with Myt1l mouse model studies, it remains to be seen whether all pathogenic mutations function by haploinsufficiency, as some could instead yield MYT1L gene products with altered functionality. We established aspects of altered neuronal development and function that may underlie patient IDD phenotypes, including upregulation of synaptic and neuronal gene expression during differentiation, followed by impaired maturation and reduced function. It would be valuable to assess additional pathogenic variants and neuronal cell types, to define both temporal and cell type-specific requirements for MYT1L for brain development and function. These findings may inform approaches for either targeting restoration of MYT1L or its key downstream targets and disrupted pathways, to define windows and approaches for treating patient phenotypes.

## EXPERIMENTAL PROCEDURES

### RESOURCE AVAILABILITY

#### Lead Contact

Further information and requests for resources and reagents will be fulfilled by the lead contact, Kristen L. Kroll.

#### Materials availability

All unique/stable reagents generated in this study are available from the lead contact with a completed Materials Transfer Agreement.

#### Data and code availability

All raw data and processed files for RNA-seq and CUT&RUN have been deposited in NCBI/Gene Expression Omnibus database (https://www.ncbi.nlm.nih.gov/geo/) as GSE244185 and GSE244189. This paper does not report original code. Any additional information required to reanalyze these data reported is available from the lead contact upon request.

### HUMAN PSC MODEL GENERATION

Proband-derived induced pluripotent stem cell models (PD-iPSC) with *MYT1L* S707Q mutation and isogenic corrected iPSC lines were generated by the Washington University Genome Engineering and Stem Cell Center (GESC). Briefly, the patient’s peripheral blood mononuclear cells were reprogrammed using CytoTune-iPS 2.0 Sendai reprogramming kit (Thermo Fisher Scientific), two iPSC heterozygous clones (PD1/2) were selected, PD clones were mutationally corrected to obtain isogenic proband derived corrected (PD-C) models, and models were Sanger sequenced to validate the mutation and correction. Lines were authenticated by STR profiling. In parallel, GESC derived human pluripotent stem cell (hPSC) models with knock-in of the same variant, by genome engineering wild-type H1 (male) human embryonic stem cells (hESC). Biomaterials were obtained from the proband with consent through Institutional Review Board (IRB) protocol 20163131. hESC work was conducted under protocol 12-002 approval by the Washington University Embryonic Stem Cell Research Oversight (ESCRO) Committee.

### STATISTICAL ANALYSES

Analyses were performed using Prism 9 software (GraphPad Software, San Diego, CA) to establish statistical significance; p<0.05 (p<0.05=*, p<0.01 =**, p<0.001 = ***). Each data point was generated from three to four independent biological replicate experiments, and acquired data were analyzed through a non-paired Student’s t-test or 2way ANOVA with Tukey’s multiple comparison’s test. Results are presented as mean +/- SEM. All original *P* values are shown in Table S9.

## ACKNOWLEDGMENTS

We thank the GESC at Washington University School of Medicine (WUSM) for generating the iPSC and VKI models, WUSM Cytogenesis & Molecular Pathology for karyotyping services, and the WU Genome Technology Access Center at McDonnell Genome Institute (GTAC@MGI) for next generation sequencing services.

## AUTHOR CONTRIBUTIONS

R.P. contributed to study design, experiments involving D30, 45 and 60 differentiation, ICC, qPCR, western blot, CUT&RUN, CRISPRi, electrophysiology (EP), puncta quantification, data analysis, and manuscript preparation. J.D. performed D60 differentiations. M.N. and R.S. performed network analysis and visualizations. M.S. conducted morphometric assays and quantification. G.C. and K.K. generated, processed, analyzed, and visualized CUT&RUN data and P.G. and B.Z. performed RNA-sequencing data processing. K.M. performed differentiation and D30 RNA preparation. B.H. generated D30, 45, and 60 confocal images. J.E.H. conducted electrophysiology experiments, analysis, and visualization and related manuscript preparation. K.L.K contributed to study design, data analysis, and manuscript preparation. All authors read and approved the final manuscript.

## DECLARATION OF INTERESTS

### Ethics approval and consent to participate

Subjects were consented for iPSC line generation by the Washington University Institutional review board of the Human Research Protection office under human studies protocol #20163131 (Dr. Kristen L. Kroll). The authors declare no competing interests.

