## Supplemental Figures S1-S8_Supplemental Figure_Table_Dataset Legends_Sup. Methods for "Autism and Intellectual Disability-Associated *MYT1L* Mutation Alters Human Cortical Interneuron Differentiation, Maturation, and Physiology"

### SUPPLEMENTAL INFORMATION

#### Figure S1

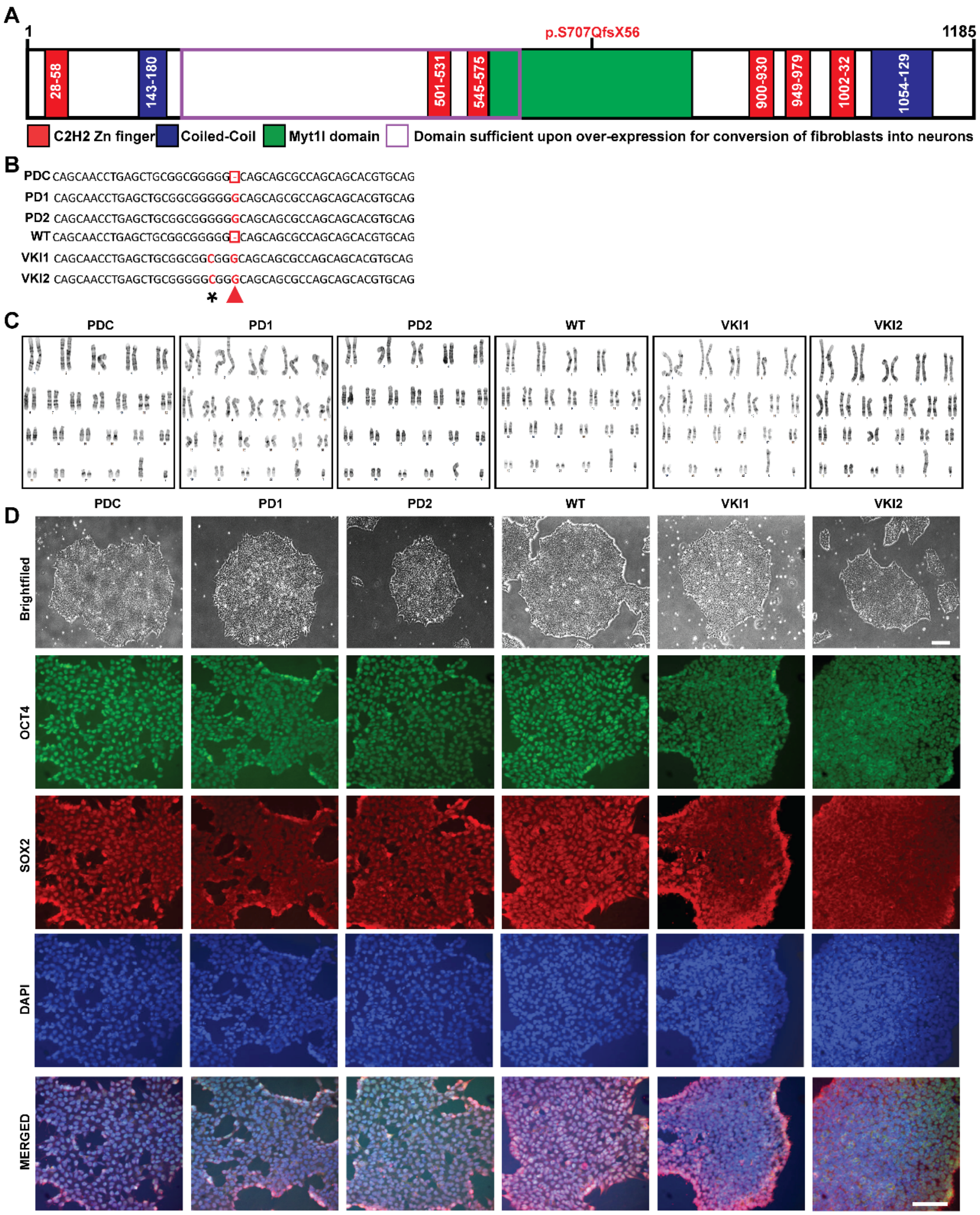

**Figure S2**

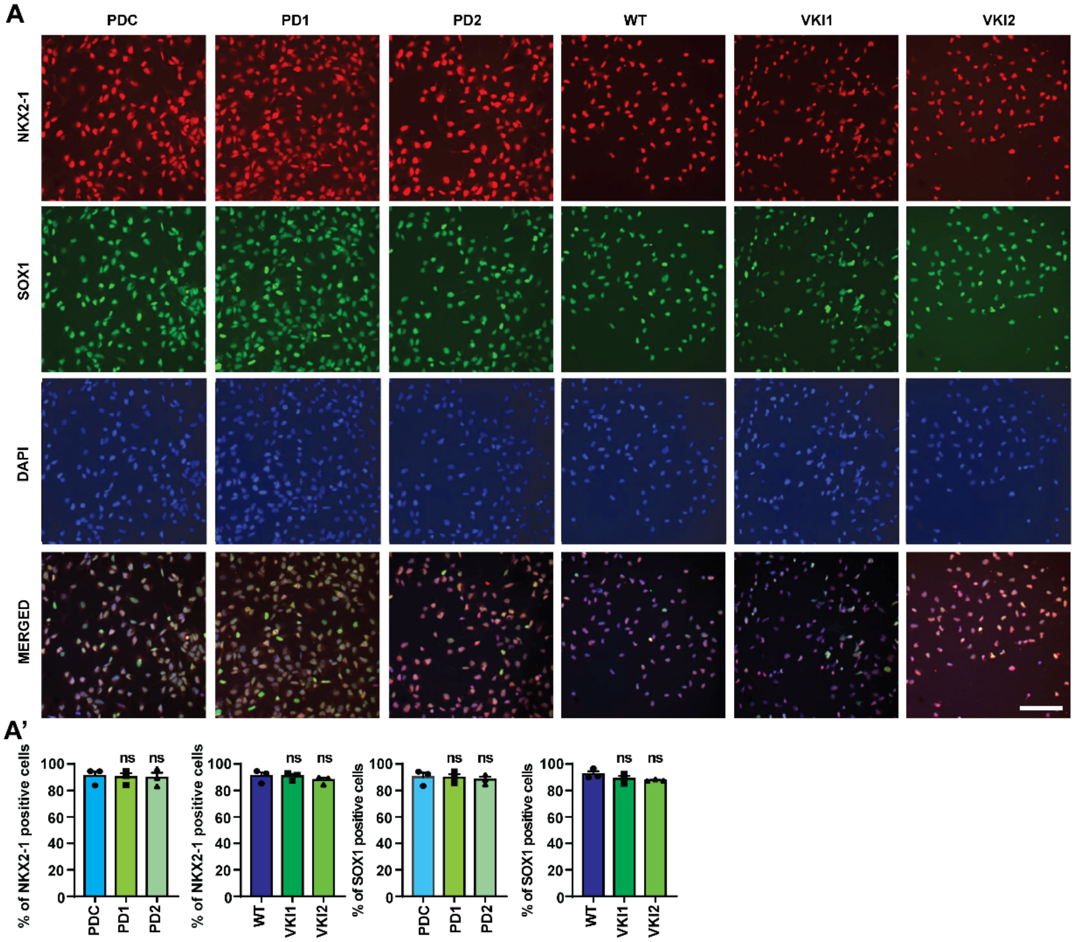
A
PDC
PD1
PD2
WT
VKI1
VKI2
NKX2-1
SOX1
DAPI
MERGED
A'
% of NKX2-1 positive cells
% of NKX2-1 positive cells
% of SOX1 positive cells
% of SOX1 positive cells
100
80
60
40
20
0
ns
PDC
PD1
PD2
WT
VKI1
VKI2

#### Figure S3

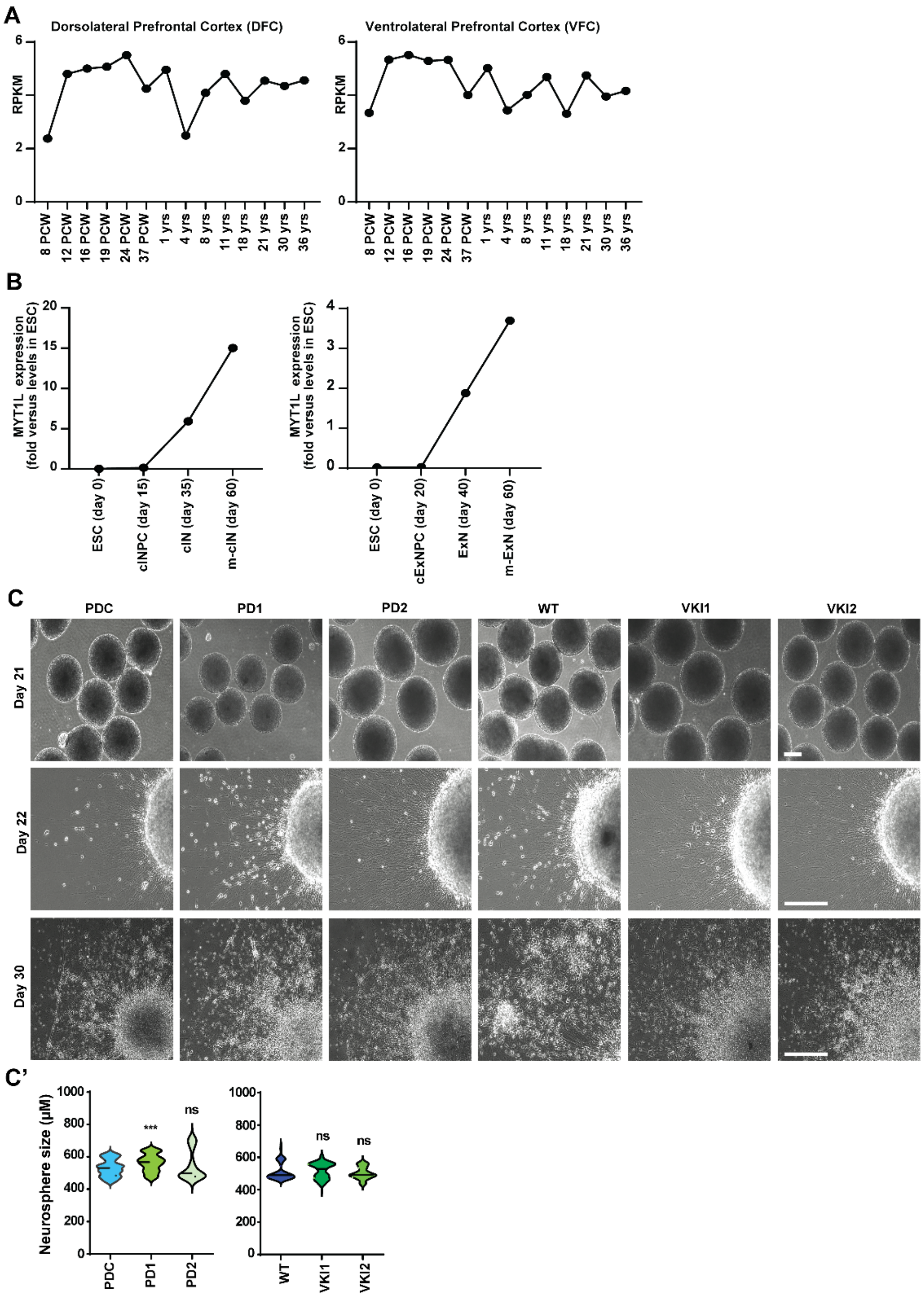

A
Dorsolateral Prefrontal Cortex (DFC)
Ventrolateral Prefrontal Cortex (VFC)
RPKM
8 PCW
12 PCW
16 PCW
19 PCW
24 PCW
37 PCW
1 yrs
4 yrs
8 yrs
11 yrs
18 yrs
21 yrs
30 yrs
36 yrs
B
MYT1L expression (fold versus levels in ESC)
ESC (day 0)
cINPC (day 15)
cIN (day 35)
m-cIN (day 60)
cExNPC (day 20)
ExN (day 40)
m-ExN (day 60)
C
PDC
PD1
PD2
WT
VKI1
VKI2
Day 21
Day 22
Day 30
C'
Neurosphere size (μM)
***
ns

**Figure S4**

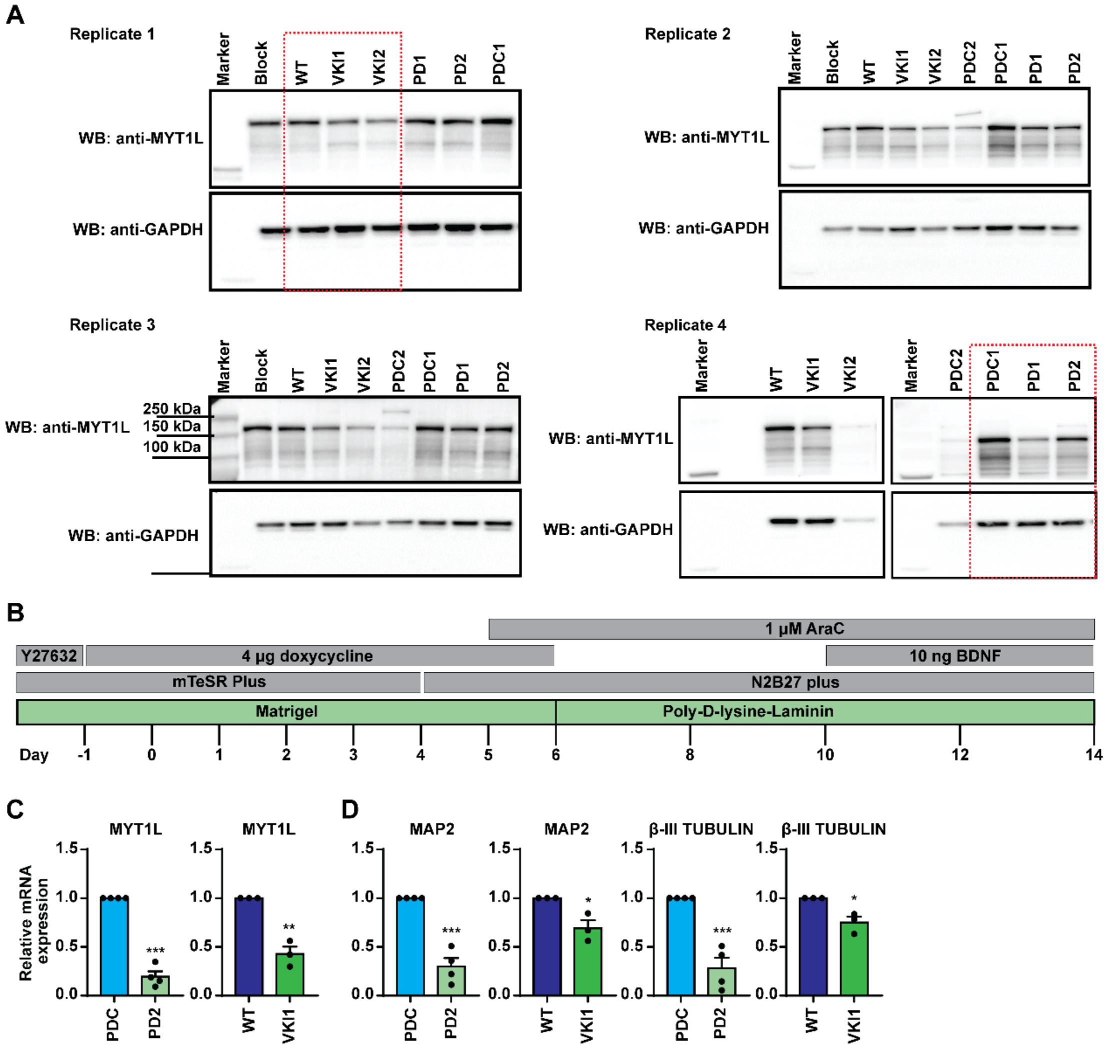

A
Replicate 1
Marker
Block
WT
VKI1
VKI2
PD1
PD2
PDC1
WB: anti-MYT1L
WB: anti-GAPDH
Replicate 2
Marker
Block
WT
VKI1
VKI2
PDC2
PDC1
PD1
PD2
WB: anti-MYT1L
WB: anti-GAPDH
Replicate 3
250 kDa
150 kDa
100 kDa
WB: anti-MYT1L
WB: anti-GAPDH
Replicate 4
WB: anti-MYT1L
WB: anti-GAPDH
B
1 μM AraC
Y27632
4 μg doxycycline
10 ng BDNF
mTeSR Plus
N2B27 plus
Matrigel
Poly-D-lysine-Laminin
Day -1 0 1 2 3 4 5 6 8 10 12 14
C
MYT1L
MYT1L
Relative mRNA expression
PDC
PD2
WT
VKI1
D
MAP2
MAP2
β-III TUBULIN
β-III TUBULIN

**Figure S5**

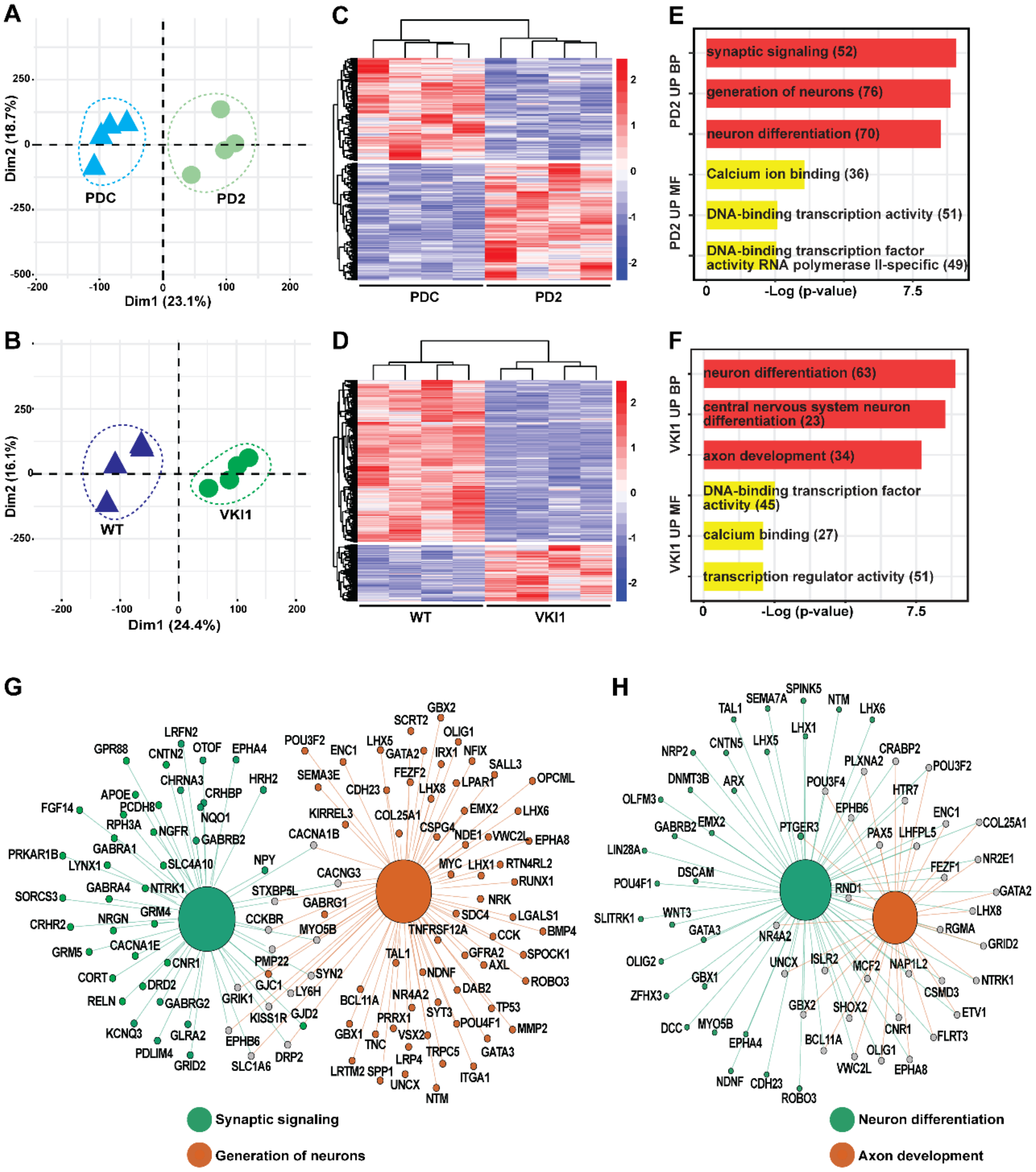
A
Dim2 (18.7%)
Dim1 (23.1%)
PDC
PD2
B
Dim2 (16.1%)
Dim1 (24.4%)
WT
VKI1
C
PDC
PD2
D
WT
VKI1
E
PD2 UP BP
PD2 UP MF
synaptic signaling (52)
generation of neurons (76)
neuron differentiation (70)
Calcium ion binding (36)
DNA-binding transcription activity (51)
DNA-binding transcription factor activity RNA polymerase II-specific (49)
-Log (p-value)
F
VKI1 UP BP
VKI1 UP MF
neuron differentiation (63)
central nervous system neuron differentiation (23)
axon development (34)
DNA-binding transcription factor activity (45)
calcium binding (27)
transcription regulator activity (51)
-Log (p-value)
G
Synaptic signaling
Generation of neurons
H
Neuron differentiation
Axon development

### Figure S6

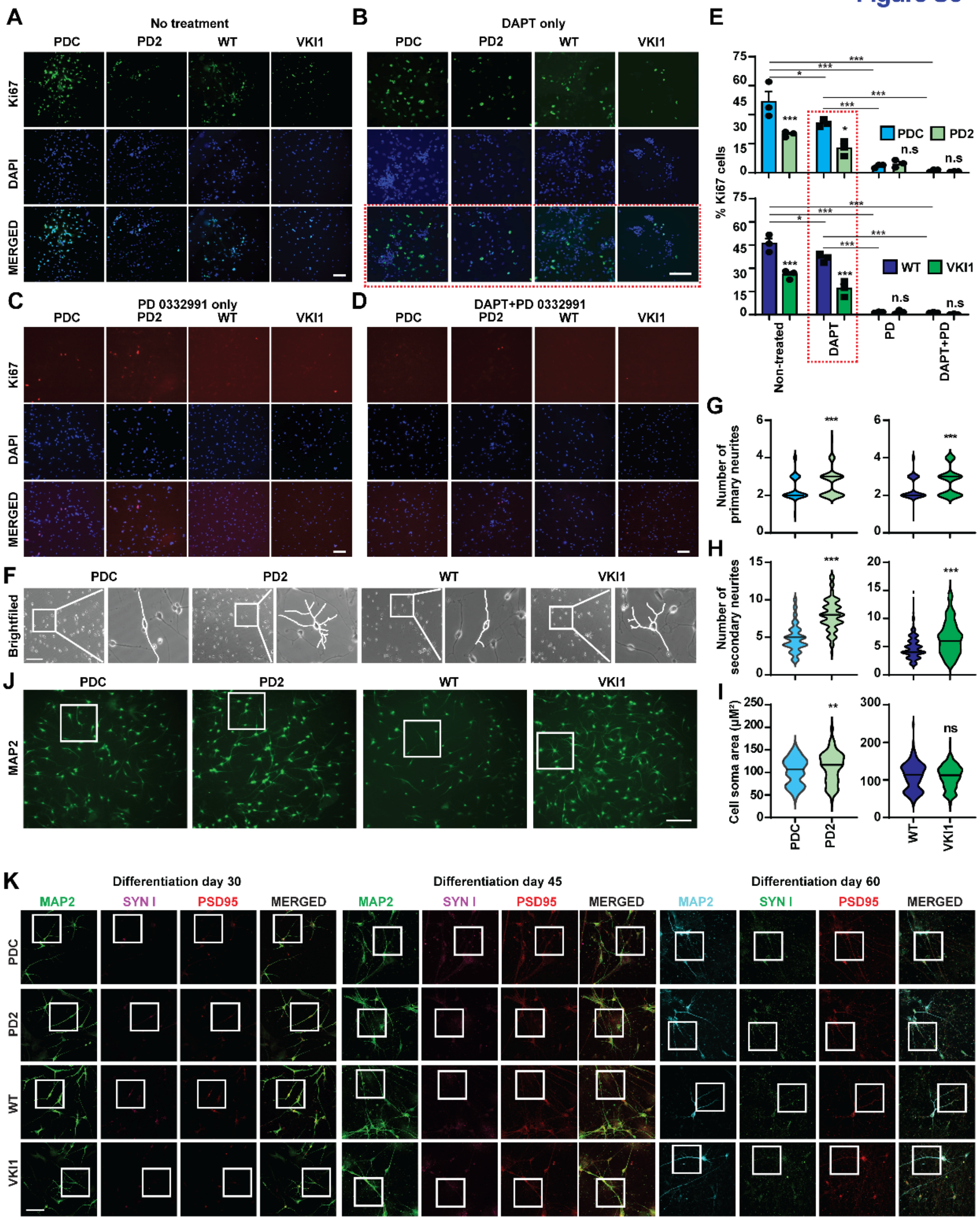
A
No treatment
PDC
PD2
WT
VKI1
Ki67
DAPI
MERGED
B
DAPT only
C
PD 0332991 only
D
DAPT+PD 0332991
E
% Ki67 cells
PDC
PD2
WT
VKI1
n.s
Non-treated
DAPT
PD
DAPT+PD
F
Brightfiled
G
Number of primary neurites
H
Number of secondary neurites
I
Cell soma area (μM²)
ns
J
MAP2
K
Differentiation day 30
Differentiation day 45
Differentiation day 60
MAP2
SYN I
PSD95
MERGED

### Figure S7

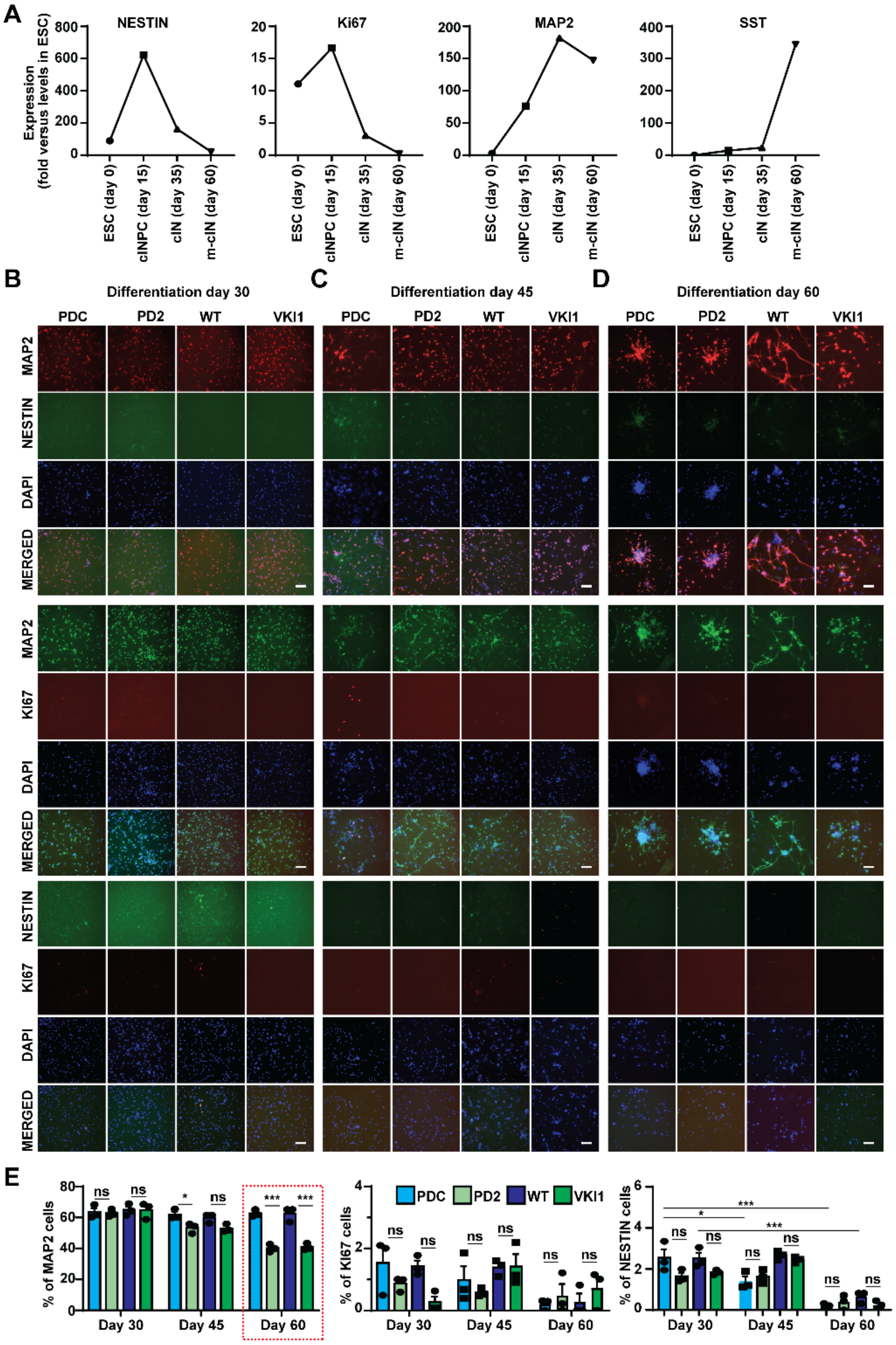
A
NESTIN
Ki67
MAP2
SST
Expression (fold versus levels in ESC)
ESC (day 0)
cINPC (day 15)
cIN (day 35)
m-cIN (day 60)
B
Differentiation day 30
C
Differentiation day 45
D
Differentiation day 60
PDC
PD2
WT
VKI1
MAP2
NESTIN
DAPI
MERGED
KI67
E
% of MAP2 cells
% of KI67 cells
% of NESTIN cells
Day 30
Day 45
Day 60
ns
*
***

#### SUPPLEMENTAL FIGURE LEGENDS

##### Figure S1. Characterization of isogenic hPSC models with and without *MYT1L* S707Q mutation

(**A**) Schematic of MYT1L protein, showing annotated domains and location of the S707Q proband variant in the MYT1L domain. MYT1L contains six zinc finger domains, two coiled-coil domains, a domain sufficient upon overexpression for conversion of fibroblasts into neurons, and a MYT1L domain of unknown function. In the proband, serine residue (S) 707 instead encodes glutamine (Q), followed by a 56 amino acid frameshift (fs) and stop codon (X). (**B**) Sequence variant present in the MYT1L cDNA in the PD1/2 and VKI1/2 models and absent in the PDC and WT lines is shown. The duplicated G is indicated in red, while the box indicates the absence of this duplication in controls. * indicates a silent mutation that was introduced to enable genome engineering to generate the VKI model hPSCs. (**C**) G-Banded Karyotyping analysis of the PDC, PD1, PD2, WT, VKI1, and VKI2 hPSC models. All lines have a normal karyotype. (**D**) Representative bright field images (taken at 4X magnification) and confocal images (taken at 20X magnification) of hPSC clones from the control (PDC, WT) and *MYT1L* S707Q variant (PD1, PD2, VKI1, VKI2) models, immunostained for the pluripotency markers OCT4 and SOX2, and counterstained with DAPI. Scale bar = 200 µM for bright field images and 100 µM for confocal images.

##### Figure S2. Characterization of PD/PDC and VKI/WT hPSC-derived cINPC

(**A**) Representative confocal images (taken at 20X magnification) of hPSC-derived cortical interneuron progenitor cells (cINPC) immunostained for NKX2.1 and SOX1, and counterstained with DAPI. Scale bar=100 µM. (**A'**). The percentage of NKX2.1 and SOX1 expressing cells, relative to all DAPI-positive nuclei. Results from three biological replicates are shown. ns: not significant.

##### Figure S3. Maturation of neurons carrying the *MYT1L* variant does not alter neurosphere size

(**A**) MYT1L expression levels in dorsolateral prefrontal cortex (DFC) (left) and ventrolateral prefrontal cortex (VFC) (right) of the human brain are shown from postconceptional weeks (PCW) 8 – 36 years. Data from Brainspan (www.brainspan.org) is expressed in reads per kilobase of transcript per million mapped reads (RPKM). (**B**) Endogenous MYT1L expression levels in stem cell (day (D) 0 hESCs), inhibitory progenitors (D15 cINPCs), immature (D30; cINs) and matured (D60; m-cINs) cortical interneurons (left), and in excitatory neuron progenitors (D20; cExNPC), and immature (D40; cExNs) and matured (D60; m-ExNs) cortical excitatory neurons (right) are shown as a fold change in expression relative to levels at day 0 (hESCs). (**C**) Representative bright-field images of neurospheres, showing neurite outgrowth in day 22 plated spheres (after plating at day 21), and in day 30 plated cINs in the models under study. Scale bar=200 µM. (**C'**) Neurosphere size was quantified from four independent biological replicates with 25 neurospheres per assay. *** $P$ <0.001 was defined by Student's t-test; ns: not significant.

##### Figure S4. *MYT1L* variant results in protein haploinsufficiency

(**A**) MYT1L protein levels shown in Figure 1C' were quantified from these uncropped western blots, with blots for the four biological replicate experiments shown. MYT1L was detected with a MYT1L-specific antibody, with GAPDH used as a loading control. The red-boxed images were cropped and are shown in Figure 1C as representative results. (**B**) Scheme for reprogramming of hPSC lines as cortical excitatory-like neurons (iNeurons) neurons by Doxycycline induced NGN2 overexpression (see Supplemental Methods for details). On day 14, mRNA expression levels were defined in NGN2 induced neurons derived from PDC, PD2, WT, and VKI1 models, with data shown derived from three to four biological replicate experiments for: (**C**) MYT1L and (**D**) the pan-neuronal markers MAP2 and β-III TUBULIN. Quantitative data are shown as the mean with +/- SEM. *P* values: *$P < 0.05$ **$P < 0.01$and ***$P < 0.001$ were defined by Student's t-test.

#### Figure S5. Transcriptomic analysis of cortical interneurons defines differential gene expression in the *MYT1L* variant PD2 and VKI1 models

(**A-B**) Principal component analysis (PCA) analysis of RNA-seq datasets encompassing four biological replicate experiments each for cINs derived from the (A) PDC versus PD2 and (B) WT versus VKI1 models. (**C-D**) Heatmap shows the expression levels of differentially expressed genes (DEGs) across four biological replicates of cINs derived from the (C) PDC-PD2 comparison and (D) WT-VKI1 comparison. Red = higher expression and blue = lower expression levels. (**E-F**) Gene ontology (GO) analysis of enriched biological processes (BP; red) and molecular functions (MF; yellow) among upregulated significantly differentially enriched genes (DEGs) in (E) PD2 and (F) VKI1 cINs, by comparison with their paired isogenic controls. (**G**) For significant DEGs upregulated in PD2 cINs, network visualization of genes related to the 'synaptic signaling' and 'generation of neurons' GO terms. Green lines and nodes are associated with 'synaptic signaling', red lines and nodes are associated with 'generation of neurons', and gray nodes are genes common to both GO terms. (**H**) For significant DEGs upregulated in the VKI1 cINs, biological process top GO terms identified gene networks associated with ‘neuron differentiation’ and ‘axon development’. In the network visualization, green lines and nodes are related to neuron differentiation, red lines are related to axon development, and gray nodes indicate genes common to both neuron differentiation and axon development.

#### Figure S6. The *MYT1L* variant reduces cell proliferation, alters neuronal morphology, and increases pre- and post-synaptic marker expression

(**A-D**) Several different conditions were utilized to initiate cINPC differentiation of the PDC, PD2, WT, and VKI1 models (see Supplemental Methods). Representative confocal images (taken at 20X magnification) are shown for cINs immunostained to detect the proliferation marker Ki67 at day 30 for: (**A**) non-treatment (no DAPT or PD 0332991), (**B**) DAPT-treatment (days 22-25), (**C**) PD 0332991 (PD)-treatment (days 22-30) or (**D**) DAPT+PD treatment (days 22-30) and counterstained with DAPI (for details see methods) Scale bar=100 μM. (**E**) Quantification of Ki67-positive cell fractions across these models under the differentiation conditions above. (**F**) Representative bright-field images (20X) of cIN morphology in the PDC, PD2, WT, and VKI models. Box panels display a magnified image corresponding to the areas shown, with tracing of the morphology of the cell soma, primary neurites, and secondary neurites. Scale bar = 100 μM. (**G-I**) The (G) number of primary neurites, (H) number of secondary neurites, and (I) quantification of the cell soma area was quantified in cINs derived from the PDC, PD2, WT, and VKI1 models. Data were derived from three biological replicate experiments, with each value derived from a total of 170-180 cINs (Table S6). (**J**) Representative microscopic images (20x) of MAP2-immunostained cINs generated from the PDC, PD2, WT, and VKI models. Boxes correspond to the enlarged images shown in **Fig. 2G**. Scale bar = 100 μM. (**K**) Representative confocal microscopic images (60X) of cINs stained with MAP2, SYN1, and PSD95 at differentiation days 30, 45, and 60 for the PDC, PD2, WT, and VKI1 models. The boxed, enlarged images and quantification is shown in **Fig. 2J-L**. Scale bar = 50 μM. Biological significance was calculated with 2way ANOVA with Tukey’s multiple comparisons test or student’s t-test. Quantitative data are shown as the mean +/- SEM. *P* values: *$P < 0.05$, **$P < 0.01$, and ***$P < 0.001$; ns: not significant. The actual *P* values are shown in Table S9.

.

**Figure S7. Neuronal differentiation was not affected by the NPC marker and proliferation marker**

**(A)** Relative expression of the NPC marker Nestin, proliferation marker Ki67, neuronal marker MAP2, and cIN marker SST in hPSCs (D0), cINPCs (D15), cINs (D35), and matured cINs (D60) are shown as fold change relative to levels at D0 (hESCs). Representative images, taken at 20X, of cINs immunostained with MAP2+NESTIN, MAP2+Ki67, or Ki67+Nestin, and counter-stained with DAPI at differentiation days 30 (**B**), 45 (**C**), and 60 (**D**) are shown for PDC, PD2, WT, and VKI models (scale bar = 100 μM). **E** shows the MAP2, NESTIN, and Ki67 positive cell percentage relative to all DAPI-positive nuclei from three biological replicate experiments. The red-boxed images were shown in Figure 5E' as representative results. Biological significance was calculated using 2way ANOVA with Tukey's multiple comparison test. Quantitative data are shown as the mean +/- SEM. *P* values: $^{*}P < 0.05$, $^{**}P < 0.01$, and $^{***}P < 0.001$. The actual *P* values are shown in Table S9.

**Figure S8. ASD and epilepsy associated genes that are MYT1L bound and significantly differentially expressed in *MYT1L* mutant cINs**

Heatmaps show expression levels (in RPKM) of ASD- (black text) and epilepsy- (red text) associated genes that are MYT1L bound and (**A**) upregulated or (**B**) downregulated DEGs in cINs derived from the PD2 and VKI1 samples versus their isogenic controls (PDC/ WT). (**C**) Summary of the findings from this study. During hPSC-cIN differentiation, *MYT1L* variant neurons (PD2, VKI1) exhibited a reduction of MYT1L expression of 30-47%, resulting in upregulation of neuronal differentiation- and synaptic signaling-related gene expression in immature cINs (D30). This effect was not observed in the CRISPRi knockdown models, which cause a stronger (68-72%) reduction of MYT1L levels. During cIN maturation (D30-60), both the *MYT1L* variant and CRISPRi models exhibit a reduction of general neuronal, mature cortical interneuron, and GABAergic neuron marker expression, supporting impaired cIN maturation. Schematic was generated in Biorender.com.

#### SUPPLEMENTAL DATASET LEGENDS

##### Dataset S1. Differential gene expression in cIN models with the *MYT1L* variant versus their isogenic controls

Differential gene expression analysis was conducted with DESeq2 as described in the Methods. Differentially expressed genes are shown for the (**A**) PDC versus PD2 (**B**) and WT versus VKI1 models in cortical interneurons (cINs), with significantly differentially expressed genes (DEGs) defined as those obtained from comparisons across four biological replicate samples in cINs that met a linear fold difference of 1.5 and adjusted p-value/FDR <0.05. (**C**) Significantly differentially expressed genes for the PDC versus PD2 comparison. (**D**) Significantly differentially expressed genes for the WT versus VKI1 comparison. (**E**) Common significantly differentially expressed genes present in both the PDC versus PD2 and WT versus VKI1 comparisons.

##### Dataset S2. Common significant DEGs for both the PDC-PD2 and WT-VKI1 comparisons

(**A-B**) RPKM values for significant DEGs for the (A) PDC-PD2 and (B) WT-VKI1 comparisons were used to generate heat maps in Figure S5C-D. **(C)** RPKM values for common significant DEGs across both the PDC versus PD2 and WT versus VKI1 comparisons were used to generate a heatmap in Figure 1D.

##### Dataset S3. Gene ontology enrichment analysis of significant DEGs in proband-derived (PD2) and variant knock-in (VKI) model cINs versus paired isogenic controls

Gene ontology terms for (**A-C**) upregulated or (**D-F**) downregulated genes. (A) up- and (D) down-regulated genes common to the PDC-PD2 and WT-VKI1 comparisons, (B) up- and (E) down-regulated genes in the PDC-PD2 comparison. (C) up- and (F) down-regulated genes in the WT-VKI1 comparison.

##### Dataset S4. Annotation and gene ontology analysis of MYT1L genome-wide occupancy

(**A**) MYT1L genome-wide occupancy in cINs was defined by CUT&RUN, with peak annotation and nearest TSS-associated gene shown, as derived from four biological replicate experiments (see Methods). (**B**) ChromHMM partitioning was used to define chromatin state at each MYT1L-bound peak. (**C**) Pathway enrichment and Gene ontology (GO) enrichment analysis of MYT1L bound peaks in cINs.

##### Dataset S5. Integration of MYT1L genome-wide occupancy with differential gene expression in *MYT1L* variant versus paired control models

(**A**) MYT1L CUT&RUN peaks were integrated with up- and down-regulated DEGs in the PDC versus PD2 or WT versus VKI1 or in both comparisons. (**B-G**) Gene ontology enrichment analysis of genes associated with MYT1L bound peaks and significantly (B) upregulated in PD2 versus PDC, (C) upregulated in the VKI1 versus WT, (D) downregulated in PD2 versus PDC, (E) downregulated in the VKI1 versus WT, (F) up-regulated in both PD2 and VKI1 models, (G) downregulated in both the PD2 and VKI1 models in cortical interneurons, by comparison with their isogenic controls.

##### Dataset S6. Integration of genes associated with MYT1L CUT&RUN peaks and significantly differentially expressed genes in the VKI and/or PD2 models, versus their isogenic controls, with ASD- and epilepsy-associated disease genes

MYT1L associated genes, derived from genome-wide occupancy (CnR) and significant DEGs were integrated with (**A**) ASD- (**B**) epilepsy-associated genes, with (**C**) indicating the total number of genes shown.

#### SUPPLEMENTAL TABLE INDEX

**Table S1.** Clinical characteristics of the *MYT1L* S707Q Proband.

**Table S2.** The clonal lines and replicates used for each panel.

**Table S3.** The list of antibodies used for immunocytochemistry (ICC) and/or western blot, with vendors and dilutions used.

**Table S4.** Primers used for RT-qPCR and CRISPRi cloning.

**Table S5.** The number of immuno-positive cells for markers assessed by ICC and the total DAPI-expressing cell counts.

**Table S6.** Quantification of numbers of primary and secondary neurites and cell soma areas.

**Table S7.** CRISPRi neurite outgrowth quantification (µm outgrowth from sphere periphery).

**Table S8.** Summary data for electrophysiology (EP).

**Table S9.** The actual *P* values and assigned *P* values used for figure legends.

#### SUPPLEMENTAL EXPERIMENTAL PROCEDURES

##### Cell lines

Human Pluripotent stem cells (hPSCs) clonal lines were grown and maintained under feeder-free conditions in mTeSR plus medium (Stemcell Technologies, #100-0274/75) on matrigel-coated plates. During standard maintenance, any hPSC colonies with morphological changes suggestive of differentiation were manually removed, hPSCs were passaged manually or using ReLeSR™ method (Stemcell technologies, #100-0483), and mycoplasma testing was regularly performed (Mycoalert Mycoplasma Detection kit (Lonza, #LT07-118)). All lines were ensured free of bacterial and fungal contamination during stock derivation, maintenance, and experimental conditions. Clonal lines were submitted to WiCell for banking as WU-PD-MYT1L-S707Qfs (PD1 clone), WU-PDC-MYT1L-CRT (PDC), and WU-VKI_MYT1LS707Qfs (VKI1). Cytogenetic analysis was performed by GTW banding, counting 20 metaphase cells and ensuring that all 20/20 cells for each clonal line (PDC, PD1, PD2, VKI1, VKI2) had a normal karyotype (Fig.S1C). hPSC and cINPC characterization and differentiation was done between passage numbers 1-7.

The Lenti-X™ 293T cell line was obtained from Takara Bio, Inc. (#632180) and maintained in growth medium (1xDMEM (Gibco, #11965-084), 10% Fetal Bovine Serum, 1% Non-Essential Amino Acids (Gibco, #11140-050), 1x sodium pyruvate (Corning, #25-000-C1), 1x GlutaMAX™ (Gibco, #35050-061), 1x Pen-Strep (Sigma-Aldrich, #P4333), 1% HEPES Buffer (Corning, #25-060-CI), 0.1% 2-mercapto ethanol (Gibco, #21985-023)). All cell lines used in this manuscript were incubated in an incubator with 5% $CO_2$ at 37°C. Detailed experimental procedures are in the Supplemental Methods.

##### hPSC specification as cortical interneuron neural progenitor cells (cINPC)

Cortical interneuron neural progenitor cell (cINPC) specification was performed as previously described (Meganathan et al. (2017); Meganathan et al. (2021); Meganathan et al. (2023)). In brief, hPSC clones were dissociated with Accutase (Gibco A11105-01) and $4x10^5$ cells/ml seeded in V-bottom 96-well plates. Embryoid bodies (EBs) were generated by plate centrifugation (200xg, 5 minutes) and incubated (37°C, 5% $CO_2$) in Neurobasal complete medium (NBM: Neurobasal A 1X, Pen/Strep 1X, GlutaMAX 1X, B27 without vitamin A 1x, Nonessential amino acid 1x, 2-Mercaptoethanol 1x) with small molecules (0.1 μM LDN193189, 10μM SB431542, 2μM XAV939, 0.1 μM SAG) for 4 days. On day 4 (D4), EBs were transferred to a non-adherent plate and incubated for 2 days at 37°C and 5% $CO_2$ on an orbital shaker. On D10, EBs were plated on Matrigel and Laminin (5μg/mL each) coated dishes. Media was replenished every other day from D6-15. On D15, cINPC EBs were dissociated with Accutase and maintained as a monolayer for up to 5 passages.

##### cINPCs differentiation into immature and mature cortical interneurons (cIN/m-cIN)

For cINPC differentiation into D30 cINs, neurospheres were made by dissociating cINPC spheres with Accutase, and $4x10^5$ cells/ml seeded in a V-bottom plate in NBM with 10μM Y-27632. Plates were incubated (37°C, 5% $CO_2$) for 4 days. On D19, Y-27632 was removed and spheres were transferred to an orbital shaker. On D21, spheres were transferred to Matrigel/laminin-coated dishes. DAPT (10μM) was included in NBM from D22–25, with cAMP (200μM), Ascorbic Acid (AA: 200μM), and BDNF (20 ng) included in NBM from D26-30. cINs were collected on day 30. To obtain D60 matured cIN (m-cIN), cINPC spheres were made as above. On D22-26, BDNF (20ng) was included in the NBM medium. On D26-32, AA (200μg), DAPT (10μM), PD0332991 (2μM), and BDNF (20ng) were included. On D32-36, AA (200μg), cAMP

(200µg), and BDNF (20ng) were included. On D36, NBM was replaced with BrainPhys$^{TM}$ Neuronal Culture Medium (Stemcell technologies #05790) with added AA (200µg), cAMP (200µg), and BDNF (20ng). GABA (300µM) and Cytarabine (1µM) were included from D40-60 to promote optimal neuronal maturation and activity. For details methods, see (Meganathan et al. (2023)). For the Ki67 immunostaining experiments that tested the consequences of different DAPT and/or PD0332991 (PD) treatment regimens for induction of cell cycle withdrawal during cIN differentiation (**Figs. S6A-E**), we added neither DAPT or PD, DAPT only on days 22-25, PD 0332991 (2µM) only on days 22-30 or PD+DAPT on days 22-30; samples were then fixed for immunocytochemistry on D30.

#### Generation of induced Neurons (iNeurons) from hPSCs

Induced neurons were generated as previously described (Schafer et al. (2019)). Briefly, a construct for doxycycline-inducible overexpression of NGN2 (Addgene #127288) was stably transduced into the PDC, PD2, WT, and VKI1 hPSC models, and selected with 0.5 µg puromycin. To induce excitatory iNeurons, hPSC models were dissociated with Accutase, and $2.5x10^5$ cells were seeded in a Matrigel-coated 6-well plate supplemented with Y-27632 (10 µM) and puromycin (0.5µg). From days 0 – 4, cells were fed with mTeSR Plus containing doxycycline (4µg) and puromycin (0.5µg). On day 4, cells were washed and media replaced with N2B27 plus (Neurobasal A (33.3%), DMEM/F-12 (66.6%), B27 supplemented with Vitamin A 1X (Gibco A35828-01), N-2 supplement (Gibco 17502-048) 1X, Pen/Strep 1X, GlutaMAX 1X, Nonessential amino acid 1X, and 2-Mercaptoethanol 1x). On day 6, the cells were dissociated with Accutase and seeded on poly-D-Lysine (10 µg/ml) and laminin (5µg/ml) coated dishes. Cytarabine (1µM) was used from D5-14. On D10, N2B27 plus was replaced with Cytarabine (1µM), 10 ng BDNF, and 5 µg Laminin, and half the medium was changed out every other day until D14. On D14, the cells were harvested and RNA was prepared as described below.

#### Morphometric analysis

At D30, images of bright field and MAP2 immunostained cINs from these models were used to measure cell soma area and quantify primary and secondary neurites. D25 cells were seeded on poly-D-Lysine (10 µg/ml) and laminin (5µg/ml) coated coverslips. Bright-field images were obtained on D30, with quantification of 170-180 individual cell soma across 3 independent biological replicate experiments counted to quantify cell soma area, and the same neurons used for primary and secondary neurite counting. MAP2-immunotained primary and secondary neurites were also counted from 120 individual cell soma across 3 independent biological replicates derived from cINs differentiated in the presence of DAPT+PD0332991. Cell soma area was quantified with ImageJ software and GraphPad Prism software used for statistical analysis. Morphometric quantifications are in Table S6. For neurite outgrowth, D21 spheres were plated on matrigel-laminin coated dishes and bright field images captured at D22. Neurite outgrowth was measured by using ImageJ software as described previously (Meganathan et al. (2021)). D22 neurite outgrowth quantification is in Table S7.

#### Immunocytochemistry (ICC) and synaptic puncta quantification

For all ICC, cells were fixed with 4% paraformaldehyde (PFA) at room temperature for 15 minutes. Then cells were blocked in 10% donkey serum, 0.1% Triton X-100, and 1% BSA in 1X phosphate-buffered saline (PBS). Primary and secondary antibodies (Key Resource Table) were diluted in dilution buffer (1% donkey serum, 0.1% Triton X-100, and 1% BSA in 1X PBS). Following blocking, cells were incubated with primary antibodies overnight at 4°C. Cells were then washed (2X; 5 minutes each) with 1X PBS and incubated for 1 hour at room temperature with secondary antibodies (donkey anti-rabbit 555 or 647, donkey anti-mouse 488, and donkey anti-goat 488, Thermo Scientific) and DAPI. Cells were washed (2X; 5 minutes each) with 1X PBS and coverslips mounted on microscope slides (Fisherbrand, #12-550-15) with ProLong™ Diamond Antifade Mount with DAPI (Invitrogen, #P36971) and allowed to dry in the dark overnight. Slides were stored at -20°C and images taken using a spinning-disk confocal microscope (Quorum) and Olympus inverted microscope with MetaMorph software or a Nikon confocal microscope with NIS-Elements imaging software or epifluorescence microscope with Micro-Manger 2.0 software.. All images were processed and quantified using ImageJ. Days 30, 45, and 60 processed images were used for synaptic puncta quantification using MATLAB-Intellicount software as described previously (Fantuzzo et al. (2017)). Images were uploaded to MATLAB-Intellicount with default thresholds with an object size 0.16 $\mu M^2$. SYN1 and PSD95 puncta were normalized to the same MAP2 area from three replicates. Antibodies and dilutions are listed in Table S3. All ICC and DAPI positive cell counts are in Table S5.

#### Reverse transcription and quantitative PCR (RT-qPCR) analysis

D30 cINs or D60 m-cINs were collected and total RNA obtained with the NucleoSpin kit (Takara, #740955). cDNA was synthesized using iScript Reverse Transcription Supermix (Bio-Rad, #1708841). Gene expression was analyzed by quantitative PCR (qPCR) using gene-specific primers with 2X qPCRBIO SyGreen Blue Mix Hi-ROX (PCRBIOSYSTEMS, #PB20.16-51). qRT-PCR data were analyzed by the $2^{-\Delta\Delta CT}$ method and normalized to RPL30 expression. qPCR primers were designed by Origene and ordered through integrated DNA technologies (IDT) or Millipore Sigma. Primers are in Table S4.

#### RNA-seq data generation and analysis

D30 cIN RNA was used for sequencing. Total RNA integrity was determined with the Agilent Bioanalyzer or 4200 Tapestation. Library preparation was performed with 500ng-1µg of total RNA. Ribosomal RNA was removed with RiboErase (Kapa Biosystems). mRNA was then fragmented in reverse transcriptase buffer and heated to 94°C for 8 minutes. mRNA was reverse transcribed to yield cDNA using SuperScript III RT enzyme (Life Technologies, per the manufacturer's instructions) and random hexamers. A second strand reaction was performed to yield double stranded cDNA. cDNA was blunt ended, an A base was added to the 3' ends, and Illumina sequencing adapters were ligated to the ends. Ligated fragments were then amplified for 12-15 cycles using primers incorporating unique dual index tags. Fragments were sequenced on an Illumina NovaSeq-6000 to obtain paired-end 150 base pair reads. RNA-seq library preparation and Illumina sequencing were performed by the Genome Technology Access Center at McDonnell Genome Institute (GTAC@MGI) at Washington University School of Medicine. An average of 30 million per reads were generated per sample. Raw reads from RNA-seq samples were processed as previously described (Meganathan et al. (2017); Chapman et al., (2024)). Gene Ontology enrichment and pathway enrichment analysis were performed with ToppGene (https://toppgene.cchmc.org/). Processed data are shown in dataset Tables S1-S3.

#### Western blotting analysis

D30 cINs were lysed in RIPA lysis buffer (Sigma, #R0278) with protease inhibitor cocktail (Thermo Scientific, #1862209) and 0.1M dithiothreitol (DTT), incubated on ice 30 minutes, and then centrifuged for 15 minutes (13,200 rpm, 4°C). Supernatant was collected and stored at -80°C or further processed for Western blotting. Protein was quantified with the Pierce™ BCA Protein Assay Kit (Thermo Scientific, #23225), and an equal amount of protein was denatured at 95°C for 5 minutes and separated on a 4-12% Bis-Tris acrylamide gel (Invitrogen, #NP0322PK2). Proteins were transferred onto 0.45 μM nitrocellulose membranes (Bio-Rad, 162-0115), blocked for 1 hour in 5% non-fat milk in PBS buffer, and membrane was incubated with specific primary antibodies (2% milk in PBS)(Key Resource Table) overnight at 4°C. HRP-conjugated secondary antibody was incubated with the membrane for 1 hour at room temperature (1:5000 dilution in 2% milk), and signals detected using an ECL kit (Thermoscientific, #34579) and the Bio-Rad Chemidoc. The antibodies and dilutions used are listed in Table S3.

#### Electrophysiology

Cultures were perfused at approximately 1 ml/min with Tyrode's solution (in mM): 150 NaCl, 4 KCl, 2 $MgCl_2$, 2 $CaCl_2$, 10 Glucose, 10 HEPES, pH adjusted to pH 7.4 with NaOH. Whole-cell electrodes had an open tip resistance of 2-6 MOhm when filled with internal solution (in mM): 140 K-glucuronate, 10 NaCl, 5 $MgCl_2$, 0.2 EGTA, 5 Na-ATP, 1 Na-GTP, and 10 HEPES, pH adjusted to 7.4 with KOH. Currents and membrane potentials were recorded with an Axopatch 200A amplifier controlled by pClamp software (Molecular Devices). Cell capacitance, input resistance, and amplitudes of inward and outward currents during depolarizing voltage steps from a holding potential of -80 mV were determined from voltage-clamp recordings (Meganathan et al. (2017)) Inactivating inward and outward currents were blocked by tetrodotoxin (TTX, 0.5 μM) and 4-aminopyridine (4-AP, 5 mM), respectively. The steady-state outward current was blocked by tetraethylammonium (TEA, 30 mM). Antagonist solutions were delivered from an 8-barrelled local perfusion pipette positioned near the recorded cell (Meganathan et al. (2021)). Current clamp recordings were obtained in a modified extracellular solution (in mM): 120 NaCl, 3 KCl, 10 glucose, 1 NaH2PO4, 4 NaHCO3, 5 HEPES, pH adjusted to 7.4 with NaOH (Meganathan et al. (2021)).

#### CRISPRi cloning and Lentivirus production

MYT1L-specific guide RNAs (gRNAs) were cloned into CROP-seq-opti (Addgene, #106280) using BsmBI restriction sites. Lentiviral expression plasmid encoding dCas9-KRAB (Addgene, #110820), and subsequently MYT1L gRNA (Key Resource Table) expression plasmids were transfected into Lenti-X™ 293T cells (Takara, #632180) using polyethyleneimine (PEI) and the packaging plasmids psPAX2 and pMD2.G (kindly provided by Prof. Andrew S. Yoo; Richner et al. (2015)). After 12 hours of transfection, growth medium (1x DMEM (Gibco, #11965-084), 10% Fetal Bovine Serum, 1% Non-Essential Amino Acids (Gibco, #11140-050), 1x sodium pyruvate (Corning, #25-000-C1), 1x GlutaMAX™ (Gibco, #35050-061), 1x Pen-Strep (Sigma-Aldrich, #P4333), 1% HEPES Buffer (Corning, #25-060-CI), 0.1% 2-mercapto ethanol (Gibco, #21985-023)) was replaced with fresh medium, collected at 48 and 72 hours post-transfection, and centrifuged at 500xg for 10 minutes. Lentivirus was concentrated with the Lenti-X concentrator (Takara, #631231) per the manufacturer's instructions, and aliquots were stored at -80°C until further use. H1 hESCs were subsequently transduced with dCas9-KRAB lentivirus and 5 μg/ml of Polybrene. After 72 hours, transduced cells were selected with 10μg/ml blasticidin. Lentivirus carrying MYT1L gRNA expression constructs was then transduced into the stable dCas9-KRAB expressing hESC line in cINPCs and selected using 1μg/ml puromycin. cINPC differentiation was performed as described above.

#### CUT&RUN Analysis

CUN&RUN was performed using the EpiCypher protocol (version 2.0). Briefly, $5x10^5$ cINs were used per assay. Cells were spun and washed twice in wash buffer (20 mM HEPES, pH 7.5, 150 mM NaCl, 0.5 mM Spermidine, 1x Roche cOmplete™ Mini EDTA-free Protease Inhibitor) and the cell pellet resuspended in 100 μl wash buffer. CUTANA™ Concanavalin A Beads (ConA beads), (EpiCypher, #21-1401) were activated in cold Bead Activation Buffer (20 mM HEPES, pH

7.9, 10 mM KCl, 1 mM $CaCl_2$, 1 mM $MnCl_2$). To bind cells to activated beads, cells were incubated with 20 µl of ConA beads for 15 minutes at room temperature. Samples were then placed on a magnet to remove supernatant and cell slurry was incubated with MYT1L (2µg) or control IgG (2µg) antibodies (Key Resource Table) prepared in antibody buffer (wash buffer, 0.01% Digitonin, 2 mM EDTA) overnight at 4°C. The next day, the cell-bead slurry was washed twice with cold digitonin buffer (wash buffer, 0.01% Digitonin), and the cell slurry was resuspended in 100µl digitonin buffer and 5µL CUTANA™ His-Tagged pAG-MNase for CUT&RUN Workflows (EpiCypher, #15-1116) was added to each reaction and incubated at room temperature for 30 minutes. The slurry was then washed twice with cold Digitonin-containing buffer. The slurry was resuspended in 50 µl digitonin buffer and targeted chromatin was released by addition of 100 mM CaCl2 for 2 hours at 4°C. Reactions were stopped by addition of stop buffer (340 mM NaCl, 20 mM EDTA, 4 mM EGTA, 50 µg/mL RNase A; Thermo Fisher Scientific, #EN0531) and 50 µg/mL Glycogen (Millipore Sigma, #10901393001), and released chromatin was purified using the CUTANA DNA purification kit (EpiCypher, #14-0050) as per the manufacturer's instructions. DNA libraries were prepared using a CUTANA CUT&RUN Library preparation kit (EpiCypher, #14-1001). Quality of the final libraries was assessed using a 4150 TapeStation System (Agilent) with High Sensitivity D1000 reagents (Agilent). Peaks from 250-750bp were combined to generate each library. Samples with unique dual-end indexes were then pooled in equimolar concentrations and sequenced by GTAC@MGI on the NovaSeq-6000 sequencer with the S4 flow cell as 2x150 bp paired-end reads to a depth of ≥10 million (M) reads per sample.

Raw data was processed using CUT&RUNTools 2.01, using de-duplicated alignment and MACS2 as the peak caller. Peak sets, including differentially bound peaks, were annotated using ChIPseeker2, defining the promotor-associated region as an area within 3 kb of the TSS. Curated peak sets were visualized with deepTools version 3, using the computeMatrix reference-point and plotHeatmap functions. Unless otherwise stated, we filtered peaks to only those within 20 kb of the nearest transcription start site (TSS) for visualization and downstream analysis. Comparisons of MYT1L binding sites to enrichment for multiple histone modifications (H3K27ac, H3K4me3, H3K27me3, and H3K9me3) was done using our previously published data (Lewis et al. (2022); Chapman et al. (2024)).

#### KEY RESOURCE TABLE

| REAGENT or | SOURCE | IDENTIFIER |
|---|---|---|
| Antibodies, See Table S3 for dilution | | |
| Rabbit monoclonal anti-Ki67 | Abcam | ab16667 |
| Rabbit polyclonal anti-Calbindin D-28K | Swant | CB-38a |
| Rabbit polyclonal anti-MYT1L | Proteintech | 25234-1-AP |
| Mouse monoclonal anti-MAP2 | Abcam | M1406 |
| Rabbit monoclonal anti-MAP2 | Cell Signaling Technology | 4542S |
| Mouse monoclonal anti-Nestin | Abcam | Ab15580 |
| Rabbit monoclonal anti-Synapsin I | Abcam | AB254349 |
| Guinea pig ant-PSD-95 PDZ domain | SYSY | 124308 |
| Rabbit monoclonal anti-Somatostatin (SST) | Invitrogen | 701935 |
| Mouse monoclonal anti-beta III Tubulin | Abcam | ab78078 |
| Polyclonal Goat anti-SOX1 | R&D systems | AF3369 |
| Rabbit polyclonal anti-SOX2 | Millipore Sigma | AB5603 |
| Mouse monoclonal anti-POU5F1 | DSHB | PCRP-POU5F1-1A4 |
| Rabbit polyclonal anti-TTF-1 | Millipore Sigma | 07-601 |
| Rabbit monoclonal anti-GAPDH | Abcam | Ab204481 |
| Rabbit polyclonal IgG | Millipore Sigma | G9545 |
| Secondary antibodies used for ICC | Thermo Fisher Scientific | https://www.thermofisher.com/antibody/secondary/query/alexa |
| Chemicals, peptides, and recombinant proteins | | |
| Y-27632 | Tocris Biosciences | 1254 |
| SB-431542 | Tocris Biosciences | 1614 |
| XAV939 | Tocris Biosciences | 374810 |
| SAG | Tocris Biosciences | 4366 |
| LDN-193189 | Stegment | 04-0074 |
| DAPT | Tocris Biosciences | 2634 |
| Ascorbic acid | Millipore Sigma | A4403 |
| BDNF | Tocris Biosciences | 2837 |
| Cytrabine | Tocris Biosciences | 4520 |
| Dibutyryl cAMP sodium | Sigma Aldrich | D0260 |

| | | |
|---|---|---|
| Palbociclib (PD0332991) | Selleckchem | S4482 |
| Critical commercial assays | | |
| CUTANA$^{TM}$ CUT&RUN Library Prep Kit | EpiCypher | 14-1001 |
| CUTANA$^{TM}$ DNA Purification Kit | EpiCypher | 14-0050 |
| Deposited data | | |
| Raw and analyzed data | This paper | GEO: GSE244185; GSE244189 |
| Experimental models: Cell lines | | |
| Human: induced Pluripotency Stem cells (iPSC)-Patient-derived MYT1L S707Q line (PD) | This paper | McDonnell Genome Institute Genome Engineering & Stem Cell Center (GESC) |
| Human: Pluripotency Stem cells Patient-derived corrected MYT1L S707Q line | This paper | hPSC Cell Line: iPSC-MYT1L-S707Q-patient-corrected clones (PDC) |
| Human: H1 hESC line | McDonnell Genome Institute Genome Engineering & Stem Cell Center (GESC)/WiCell | hESC Cell Line: H1 |
| Human: H1 hESC MYT1LS707Q variant knock-in clone | This paper | hESC Cell Line: H1-MYT1L-S707Q-VKI-clones (VKI-clones) |
| LentiX-293T | Takara | 632180 |
| Oligonucleotides, See Table S4 for all primers used in this manuscript. | | |
| Guide RNA: MYT1L-G1-Forward | Sanson et al. (2018) | 5'-GCAGTCTTGTTGAAGCACCC-3' |
| Guide RNA: MYT1L-G1-Reverse | | 5'-GGGTGCTTCAACAAGACTGC-3' |
| Guide RNA: MYT1L-G1-Forward | Sanson et al. (2018) | 5'-GGTGCTTCAACAAGACTGCA-3' |
| Guide RNA: MYT1L-G1-Reverse | | 5'-TGCAGTCTTGTTGAAGCACC-3' |
| Quantitative PCR: β-III TUBULIN-Forward | Tang et al. (2020) | 5'-TCAGCGTCTACTACAACGAGGC-3' |
| Quantitative PCR: β-III TUBULIN -Reverse | | 5'-GCCTGAAGAGATGTCCAAAGGC-3' |
| Quantitative PCR: DLX2-Forward | Meganathan et al. (2017) | 5'-TACTCCGCCAAGAGCAGCTATG-3' |
| Quantitative PCR: DLX2-Reverse | | 5'-CGAATTTCAGGCTCAAGGTCCTC-3' |
| Quantitative PCR: GABRB2-Forward | Sanson et al. (2018) | 5'-GGGATGAACATTGACATTGCCA-3' |
| Quantitative PCR: GABRB2-Reverse | | 5'-CTCCAGGCTTGTTGAAAGTACAT-3' |

| | | |
|---|---|---|
| Quantitative PCR: GAD2-Forward | Meganathan et al. (2017) | 5'-GCCAACTCTGTGACGTGGAATC-3' |
| Quantitative PCR: GAD2-Reverse | | 5'-GCTGAAAGAGGTAGGAGGCATG-3' |
| Quantitative PCR: Ki67-Forward | Sanson et al. (2018) | 5'-TCCTTTGGTGGGCACCTAAGACCTG-3' |
| Quantitative PCR: Ki67-Reverse | | 5'-TGATGGTTGAGGTCGTTCCTTGATG-3' |
| Quantitative PCR: LHX1-Forward | Tang et al. (2020) | 5'-GCCAAAGAGAACAGCCTTCACTC-3' |
| Quantitative PCR: LHX1-Reverse | | 5'-GGTCGTCATTCTCGTTGCTACC-3' |
| Quantitative PCR: MAP2-Forward | Tang et al. (2020) | 5'-AGGCTGTAGCAGTCCTGAAAGG-3' |
| Quantitative PCR: MAP2-Reverse | | 5'-CTTCCTCCACTGTGACAGTCTG-3' |
| Quantitative PCR: NKX2-1-Forward | Li et al. (2012) | 5'-AGGACACCATGAGGAACAGC-3' |
| Quantitative PCR: NKX2-1-Reverse | | 5'-GCCATGTTCTTGCTCACGTC-3' |
| Commercial Plasmid DNA | | |
| Lenti-dCas9-KRAB-blast | Xie et al. (2017) | Addgene Plasmid #89567 |
| CROP-seq-opti | Hill et al. (2018) | Addgene Plasmid #106280 |
| Software and algorithms | | |
| STAR Version 2.5.4b | Dobin et al. (2013) | https://github.com/alexdobin/STAR |
| FeatureCounts Version 1.6.4 | Liao et al. (2013) | |
| DESeq2 Version 3.18 | Love et al. (2014) | https://bioconductor.org/packages/release/bioc/html/DESeq2.html |
| ChIPSeeker Version 1.32.1 | Yu et al. (2015) | https://github.com/YuLab-SMU/ChIPseeker |
| DeepTools Version 3.5.4 | Ramirez et al. (2016) | https://github.com/deeptools/deepTools |
| ImageJ | Schneider et al. (2012) | https://imagej.nih.gov/ij/ |
| ToppFun Tool | Chen et al. (2007) | https://toppgene.cchmc.org/help/publications.jsp |
| GraphPad Prism Version 9 | GraphPad Software | https://www.graphpad.com |
| Rstudio Version 3.5.1 with R Version 4.2.1 | Rstudio Software | https://www.rstudio.org |
| ImageJ | Schneider et al. (2012) | https://imagej.nih.gov/ij/ |
| MetaMorph | MetaMorph Software | https://www.moleculardevices.com/products/cellular-imaging-systems/acquisition-andanalysis-software/metamorph-microscopy |
| Chemidoc Image Lab Touch Software | Bio-Rad | https://www.bio-rad.com/enus/ product/chemidoc-mp-imagingsystem?ID=NINJ8ZE8Z |

| | | |
|---|---|---|
| Intellicount | Fantuzzo et al. (2017) | https://techfinder.rutgers.edu/tech/Intellicount |
| StepOnePlus™ Real-Time PCR System with StepOne Software Version 2.3 | Applied Biosystems | https://www.thermofisher.com/order/catalog/product/4376600 |
| CUT&RUN Tools 2.0 | Yu et al. (2021) | https://pubmed.ncbi.nlm.nih.gov/34244724/ |
| Other | | |
| BrainSpan MYT1L expression levels in RPKM for 8 PCW – 36 years | BrainSpan Atlas of the Developing Human Brain | https://www.brainspan.org/ |
